# Adaptive introgression of a visual preference gene

**DOI:** 10.1101/2023.07.12.548653

**Authors:** Matteo Rossi, Alexander E. Hausmann, Pepe Alcami, Markus Moest, Daniel Shane Wright, Chi-Yun Kuo, Daniela Lozano, Arif Maulana, Lina Melo-Flórez, Geraldine Rueda- Muñoz, Saoirse McMahon, Mauricio Linares, W. Owen McMillan, Carolina Pardo-Diaz, Camilo Salazar, Richard M. Merrill

## Abstract

Visual preferences are important drivers of mate choice and sexual selection, but little is known of how they evolve at the genetic level. Here we take advantage of the diversity of bright warning patterns displayed by *Heliconius* butterflies, which are also used during mate choice. We show that two *Heliconius* species have evolved the same visual mating preferences for females with red patterns by exchanging genetic material through hybridization. Extensive behavioral experiments reveal that male preferences are associated with a genomic region of increased admixture between these two species. Variation in neural expression of *regucalcin1*, located within this introgressed region, correlates with visual preference across populations, and disruption of *regucalcin1* with CRISPR/Cas9 impairs courtship towards conspecific females, proving a direct link between gene and behavior. Our results support a role for hybridization during behavioral evolution, and show how visually-guided behaviors contributing to adaptation and speciation are encoded within the genome.

---

Organisms often use color, and other visual cues, to attract and recognize suitable mates (*1*). The evolution of these cues is increasingly understood at the molecular level, providing insights into the nature and origin of genetic variation on which selection acts *e.g.*, (*2–7*). However, we know little of the genetic mechanisms underlying variation in the corresponding preferences, or visually guided behaviors more broadly. Indeed, while progress has been made for other sensory modalities (and especially chemosensation, *e.g.*, (*8–10*)), genetic studies of visual preference evolution remain limited to the identification of relatively broad genomic regions containing tens or hundreds of genes, and/or are unable to distinguish between causal and correlated genetic changes (*11–15*). Although these studies have undoubtedly contributed to our understanding of population divergence, identifying the causal genes involved is key to uncovering how behavioral variation is generated during development and across evolutionary time.

*Heliconius* butterflies are well known for their diversity of bright warning patterns, which are also used as mating cues (*16*). Closely related taxa often display divergent wing patterns, and because males almost invariably prefer to court females that share their own color pattern, this contributes an important premating reproductive barrier between species *e.g.*, (*17*). While the genetics and evolutionary history of *Heliconius* color pattern variation is well understood (*18–24*), we know very little of the specific genetic mechanisms contributing to the evolution of the corresponding visual preference behaviors. Previously we identified three genomic regions controlling differences in male courtship behaviors between the closely-related sympatric species *H. cydno* and *H. melpomene*, which differ in color pattern (*11*). However, further fine mapping of this behavioral phenotype is impractical, and even the best supported of these behavioral quantitative trait loci (QTLs), which has also been explicitly linked to differences in visual preference (*25*), is associated with a confidence region containing 200 genes. Although patterns of neural gene expression highlight a number of candidates (*26*), the exact genes involved remain unknown.

Here we take advantage of the mimicry relations among three closely related *Heliconius* species to determine how genetic variation for visual preferences has evolved in relation to that of the corresponding color pattern cues. Whereas west of the Eastern Cordillera in the Andes coexisting *H. cydno* and *H. melpomene* differ in forewing color (being white and red respectively), on the eastern slopes *H. cydno* is replaced by its sister species *H. timareta*, which shares the red patterns of the local *H. melpomene* (Fig. 1A). Mimicry between these two red species is not the result of independent mutations, but adaptive introgression, whereby *H. timareta* acquired color pattern alleles following hybridization with *H. melpomene* (*23, 24, 27*). This presents an excellent opportunity to both i) test whether behavioral phenotypes can similarly evolve through the reassembly of existing genetic variants on a novel genomic background, and ii) to isolate the causal genes. We identify a region of increased admixture between *H. melpomene* and *H. timareta* that is strongly associated with parallel preferences for red females in both species. We then leverage this finding alongside transcriptomic analysis and genome-editing to identify a major effect gene underlying the evolution of visual preferences.

**Figure 1.**
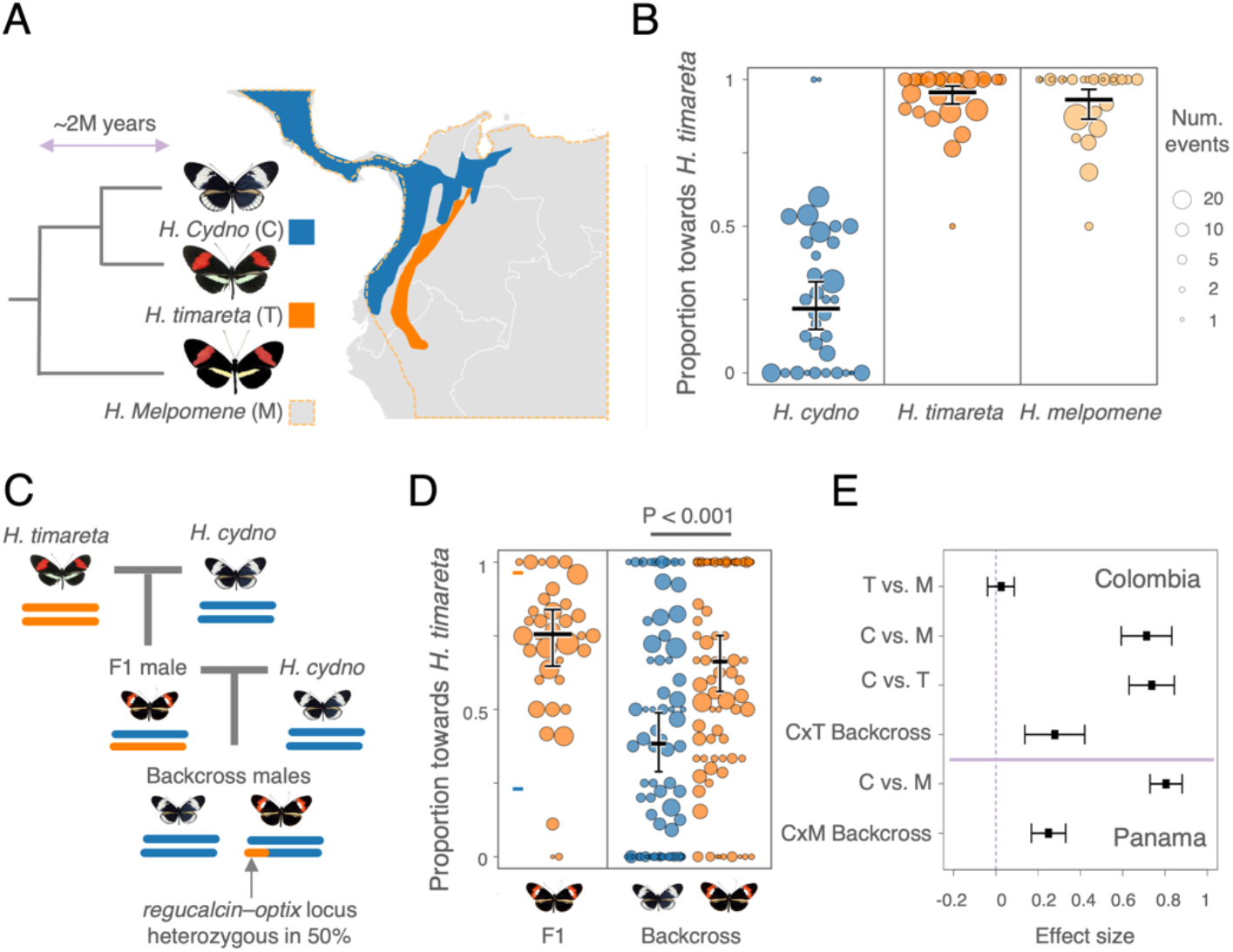
Parallel visual preferences are controlled by the same genomic region in the *Heliconius melpomene-cydno* group. (**A**) *H. melpomene* (dotted orange line) co-occurs with *H. cydno* (blue) in Central America and South America to the west of the Eastern Cordillera in the Andes, while *H. melpomene* co-occurs with *H. timareta* (orange) to the east of the Eastern Cordillera. *H. melpomene* and *H. timareta* share red warning patterns even though the latter is more closely related to the white/yellow *H. cydno*. (**B**) Proportion of courtship time directed towards red *H. timareta* females relative to white *H. cydno* females by males of the three species. Point size is scaled to the number of total minutes a male responded to either female type (a custom swarmplot was used to distribute dots horizontally). Estimated marginal means and their 95% confidence intervals are displayed with black bars. (**C**) Crossing design for producing backcross hybrid individuals to *H. cydno* segregating at the behavioral QTL region on chromosome 18. (**D**) Relative courtship time directed towards red *H. timareta* females by F1 hybrid and backcross to *H. cydno* hybrid males. Orange points represent individuals that are heterozygous (*i.e.*, ‘cydno-timareta’) and blue points represent individuals that are homozygous for *H. cydno* alleles at the QTL peak/*optix* region on chromosome 18. (**E**) Differences in estimated marginal means for relative courtship time between butterfly types tested in Colombia (this study) and in Panama (*9*). T= *H. timareta*, M= *H. melpomene*, C= *H. cydno,* Backcross *=* backcross to *H. cydno* hybrids.

## Evolution of parallel visual preference behaviors

To explore the evolution of visually guided behaviors across the *melpomene*-*cydno* group we assayed mate preference for populations sampled across Colombia. Specifically, we tested *H. melpomene* and *H. timareta* males from the eastern slopes of the Eastern Cordillera, which both have a red forewing band, as well as *H. cydno* males from the western slopes of the Eastern Cordillera, which have a white or yellow forewing band. Male butterflies were simultaneously presented with a red *H. timareta* and a white *H. cydno* female in standardized trials. Males of the two red species showed a stronger preference for red females than the *H. cydno* males (differences in proportion courtship time towards red females: *H. timareta* - *H. cydno* = 0.737 [0.630 – 0.844], *H. melpomene* - *H. cydno* = 0.713 [0.593 – 0.832]; *n* = 87, 2Δln*L* = 99.8, *P* << 0.001; Fig. 1B), but there was no difference in mate preference between the two red species (0.025 [-0.039 – 0.087]). We confirmed that preference differences between male *H. timareta* and *H. cydno* are largely based on visual cues by repeating our experiment, this time presenting males with two *H. cydno* females, where the forewings of one were artificially colored to match the red forewing of *H. timareta* (with respect to *Heliconius* color vision), and the wings of the other with a transparent marker as a control (*H. timareta* – *H. cydno* = 0.46 [0.36 – 0.56]; *n* = 94, 2Δln*L* = 53.7, *P* << 0.001, Fig S1). Overall, these results closely mirror previous data for Panamanian populations of *H. cydno* and *H. melpomene* (*11, 17*), where the latter shows a much stronger preference for red females, and confirms that although *H. timareta* is more closely related to *H. cydno*, it shares the visual preference phenotype of *H. melpomene*.

## The same major effect locus contributes to red preference in *H. melpomene* and *H. timareta*

If introgression has contributed to this parallel behavioral evolution for females with red patterns, we would expect the same genomic locations to influence the preference behaviors of both *H. melpomene* and *H. timareta*. In other words, we expect that the alleles at the location of the *H. melpomene* x *H. cydno* QTLs also segregate with preference differences in crosses between *H. timareta* and *H. cydno*. Confirming this, we found that genotype at the end of chromosome 18 is a strong predictor of male preference in *H. timareta* x *H. cydno* hybrids. Specifically, backcross hybrid males that inherit an allele from *H. timareta* at the previously detected QTL peak spent more time courting red *H. timareta* than white *H. cydno* females, compared to their brothers that inherited two copies of the *H. cydno* alleles at the same location (differences in proportion courtship time between males with ‘cydno-timareta’ and ‘cydno-cydno’ genotypes *=* 0.279 [0.137 – 0.42]; *n* = 157, 2Δln*L* = 14.02, *P* = 0.00018; Figs. 1C and D). Notably the effect size observed here is almost identical to that seen in hybrids between *H. cydno* and *H. melpomene* (*i.e.* 0.249 [0.168 – 0.33]; Fig. 1E).

To further confirm that the QTL region on chromosome 18 specifically modulates visual mate preferences, we also assayed mate preference behaviors of *H. timareta* x *H. cydno* hybrid males towards white (transparently-painted) and red-painted *H. cydno* females (as described above). We found that backcross males heterozygous for *H. timareta* and *H. cydno* alleles at QTL confidence region on chromosome 18 court red-painted females more frequently than their brothers homozygous for the *H. cydno* allele (*n* = 270, 2Δln*L* = 7.811, *P* = 0.005, Fig S1). While the effect size for this experiment (0.0778 [0.024 – 0.13]) is reduced compared to that seen for experiments using *H. timareta* females, this still represents a considerable proportion of the observed parental difference (∼17%). Together our two experiments confirm that the same genomic region at the end of chromosome 18 modulates variation in visual mate preferences across the *melpomene*-*cydno* group.

## Genomic signatures of adaptive introgression at the preference locus

To further determine whether introgression of preference alleles has contributed to behavioral evolution in these species, we next analyzed admixture proportions (*f*_d,_ (*28*)) between sympatric red-preferring *H. melpomene* and *H. timareta.* We observed two striking peaks of admixture in the QTL region on chromosome 18, located within the behavioral QTL peak (*i.e.* the region of greatest statistical association with difference in male preference between *H. cydno* and *H. melpomene*) and upstream of the adjacent major color pattern gene, *optix* (corresponding to its putative regulatory region) (Fig. 2, *c.f.* Fig. S2). Admixture estimates are repeatable across geographic populations of *H. melpomene* and *H. timareta*, and are independent of variation in local recombination rates (known to otherwise correlate with admixture proportions (*29*) (Fig. 2).

**Figure 2.**
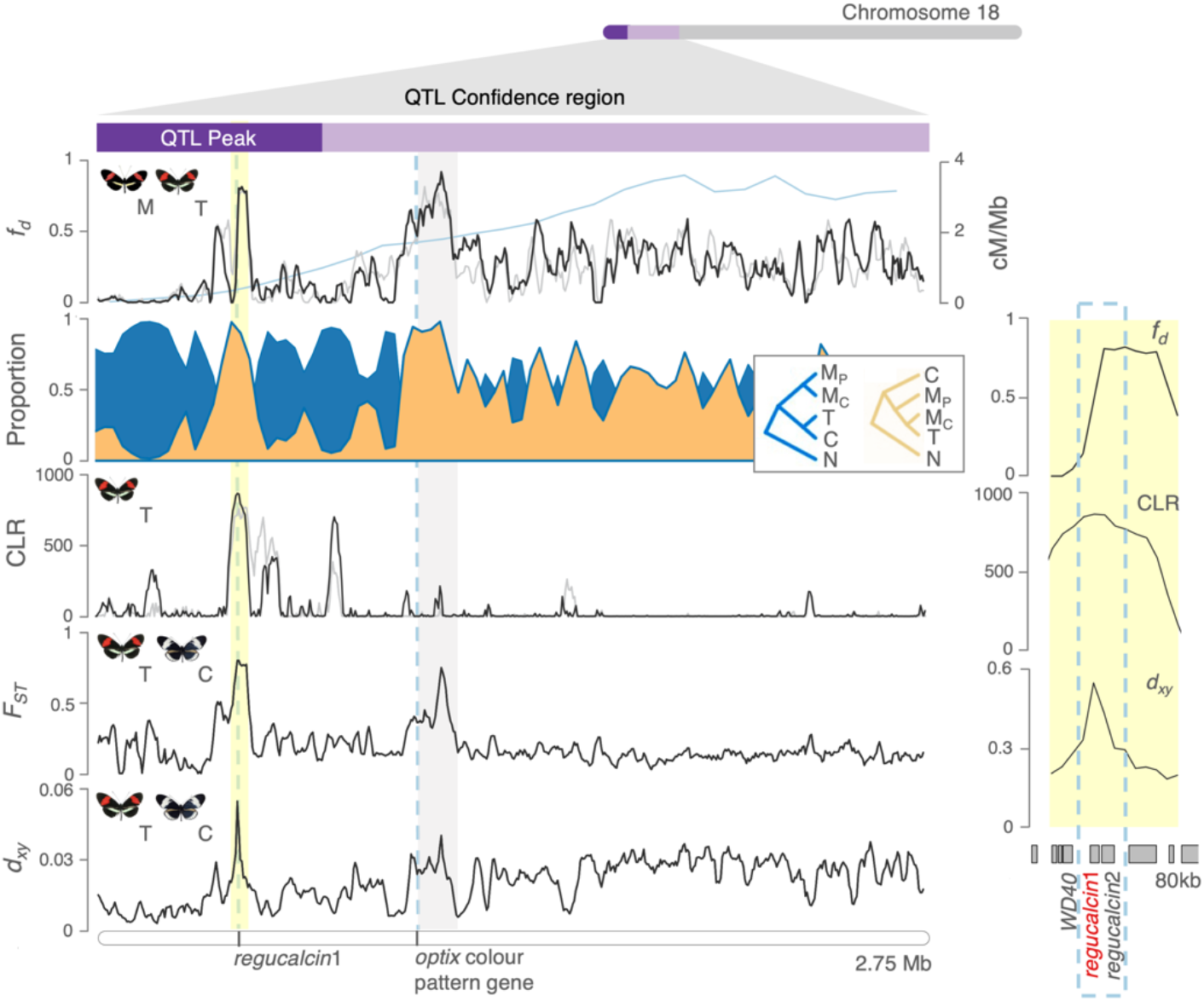
Different genomic signatures support both divergence and adaptive introgression at the *regucalcin* locus. Left, from top to bottom: Admixture proportion values (20kb windows) between *H. melpomene* and *H. timareta* at the behavioral QTL region on chromosome 18 (x-axis indicates physical position) for Colombian (black) and Peruvian (gray) populations, with recombination rate overlaid in blue; topology weightings (proportions of a particular phylogenetic tree over all possible rooted trees) for the “species” (blue) and “introgression” (orange) trees (50 SNPs windows, a *loess* smoothing function across 150kb windows was applied). *H. numata* was used as outgroup; composite likelihood ratio (CLR) of a selective sweep in *H. timareta* (50 SNPs windows); fixation index (F_ST_) and d_xy_, measures of genetic differentiation and divergence, between *H. timareta and H. cydno*. The gene coordinates of the candidate gene for behavioral difference *regucalcin1* as well as *optix* (∼500 kb apart) and its putative regulatory regions, are highlighted by vertical gray dotted lines and shading. Panel to the right zooms into the region containing candidate behavioral genes. M, T, C and N denote *H. melpomene*, *H. timareta, H.cydno* and *H. numata,* respectively; subscripts _P_ and _C_ denote Panama and Colombia, respectively

Introgression at the two loci on chromosome 18 is further supported by analyses using *Twisst* (*30*), which quantifies the proportion of different phylogenetic relationships among individuals of different species across the chromosome. In these analyses, the “introgression” topology, where *H. timareta* and *H. melpomene* cluster together, with *H. cydno* as an outgroup, is strongly supported both within the QTL peak and at *optix* (Figs. 2 and S3). These admixture peaks additionally coincide with elevated levels of genetic differentiation (*F*_ST_) and absolute genetic divergence (d_xy_) between red- and white-preferring populations (Fig 2). Finally, using *Sweepfinder2* (*31*), we found evidence for a recent selective sweep in *H. timareta* (top 1% quantile across autosomes), coincident with the peak of increased admixture within the behavioral QTL peak described above, but not at *optix* (Figs. 2 and S4). These results suggest adaptive introgression of alleles from red-preferring *H. melpomene* into *H. timareta* at a genomic location strongly associated with variation in visual preference.

## *Cis-*regulated expression differences of *regucalcin1* are associated with visual preference

We next generated RNAseq libraries for combined eye and brain tissue from adult males across all populations tested in our preference assays to determine whether consistent differences in gene expression are associated with the behavioral QTL on chromosome 18. We sampled at the adult stage reasoning that if the neural mechanism underlying divergent behaviors involves a change in neuronal activity, this might require sustained transcription. Of 200 genes within the chromosome 18 QTL candidate region, only one was consistently differentially expressed across all red and white preferring population comparisons (reared under common garden conditions, Fig. S5). Specifically, *regucalcin1*, which perfectly coincides with the peak of adaptive introgression between red-preferring populations detected above, shows lower expression in the neural tissue of Panamanian and Colombian populations of *H. melpomene* and *H. timareta*, all of which we have shown to have a red preference as compared to *H. cydno* (Fig. 3A and S6). Expression of *regucalcin1* is also significantly reduced in *H. melpomene amaryllis* and *H. melpomene melpomene* as compared to *H. cydno*, two additional populations additionally known to display a preference for red females (*17, 32*) (Fig. S6). Immunostainings in adult male *H. melpomene* brains revealed expression patterns of *regucalcin1* across the brain, including in the visual pathways, predominantly in soma, and also in neuropil (Fig. 3C). Although this does not pinpoint the particular site of action in the brain, it confirms that regulatory changes of *regucalcin1* have the potential to affect visual preference behavior.

**Figure 3.**
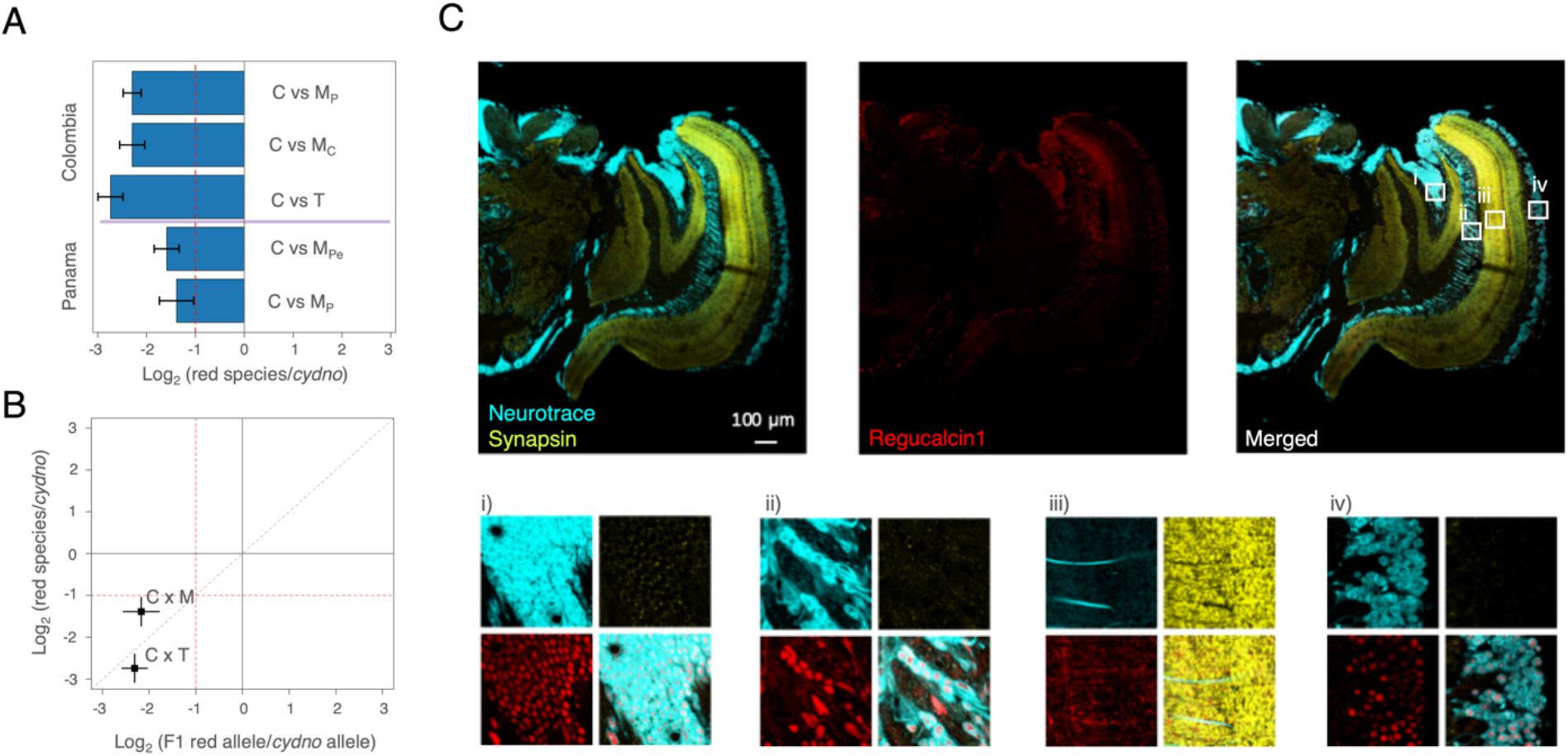
*Cis-*regulated expression differences of *regucalcin1* are associated with visual preference and *regucalcin1* is expressed in the visual pathways. (**A**) *Regucalcin1* is differentially expressed between red-preferring and white-preferring butterflies. Histogram heights represent the value and bars the standard error of the (base 2) logarithmic fold change in expression between red-preferring and white-preferring *Heliconius* subspecies (comparisons conducted only between butterflies raised in the same insectary locations). The dashed red line indicates the threshold for a 2-fold change in expression. M, T, C denote *H. melpomene*, *H. timareta* and *H.cydno,* respectively; subscripts _P, C_ and _Pe_ denote Panama, Colombia and Peru, respectively. (**B**) Allele specific expression analyses indicate that differences in expression of *regucalcin1* in the brains of red and white preferring population is *cis*-regulated. Points indicate the value and bars the standard error of the log2 (fold change) in expression between parental species (vertical) and the alleles in F1 hybrids (horizontal), for *regucalcin1*. Dashed red lines indicate the threshold for a 2-fold change in expression for the genes in the species (horizontal), and for the alleles in the hybrids (vertical). *Regucalcin1* is largely *cis*-regulated (indicated by proximity to y=x). (**C**) *Regucalcin1* is widely expressed in *Heliconius melpomene* brains, including the visual pathway. On top, immunostaining of the right hemisphere, from left to right: counterstaining of somata with *neurotrace* and of the neuropil with *synapsin*, center: staining against *regucalcin1*, right: merged image. Below, enlargement of somata (**i, iii, iv)** and neuropil **(ii)** along the visual pathway.

If expression differences in *regucalcin1* are responsible for the behavioral variation associated with the QTL on chromosome 18, they must result from changes within the *cis*-regulatory regions of the genes themselves, as opposed to those of other *trans*-acting genes elsewhere in the genome. To test whether differences in gene-expression levels between parental species were due to *cis*- or *trans*-regulatory changes, we conducted allele-specific expression analyses in adult male F1 *H. melpomene* x *H. cydno* and *H. cydno* x *H. timareta* hybrids. In F1 hybrids, both parental alleles are exposed to the same *trans*-environment, and consequently *trans*-acting factors will act on alleles derived from each species equally (unless there is a change in the *cis*-regulatory regions of the respective alleles). Confirming *cis-*regulation of *regucalcin1*, we found a significant 2-fold up-regulation of the *H. cydno* allele relative to the *H. melpomene* or *H. timareta* allele in the neural tissue of both our *H. melpomene* x *H. cydno* and *H. timareta* x *H. cydno* F1 males (Wald test all comparisons: *P* < 0.001, Fig. 3B).

## CRISPR/Cas9 mediated knock-out of *regucalcin1* disrupts male courtship behaviors

Combining genetic crosses and behavioral data, as well as population genomic and expression analyses, our results strongly implicate *regucalcin1* as a visual preference gene. To functionally test for a link between gene and behavior, we knocked-out the protein coding region of *regucalcin1* in *H. melpomene* individuals by introducing a deletion spanning most of its first and second exon using CRISPR/Cas9 (Fig. 4A). In trials with a single conspecific female (Fig. 4B), mosaic knock-out (mKO) males (*i.e.*, those with a deletion at *regucalcin1* in a substantial number of cells, including in brain tissue, Fig. S8) were significantly less likely court than control (ND) males without the deletion (difference in proportion minutes courting, trials with mKO males - trials with ND males = 0.24 [0.03-0.55]; 2*ΔlnL* = 4.51, *P* < 0.05; Fig. 4C). mKO knockout individuals may suffer decreased viability both pre- and post-eclosion (Fig. S7), and some mKO butterflies were unable to fly (8/44 individuals) as determined in our ‘drop test’, as compared to 0/40 ND individuals or 0/42 wildtype individuals; Fisher exact test: *P* < 0.001). However, only surviving males that could fly were included in our courtship trials. Furthermore, all mKO (36/36), ND (31/31) and wildtype (30/30) individuals tested, including seven individuals that failed the subsequent drop test, showed an optomotor response, suggesting basic visual sensorimotor skills are largely intact in mKO individuals. Finally, we observed no difference in the proportion of time flying or feeding between the same mKO or ND males included in our courtship trials (0.01 [-0.07-0.097]; 2*ΔlnL* = 0, *P* > 0.9; Fig. 4C and S9). In other words, courtship – but not other behaviors – was significantly reduced in *regucalcin1* knockout males as compared to controls, which retain functional copies of *regucalcin1*. This provides functional evidence that *regucalcin1* has a specific effect on male courtship behavior, and that this is not due to a more general impairment of behavior.

**Figure 4.**
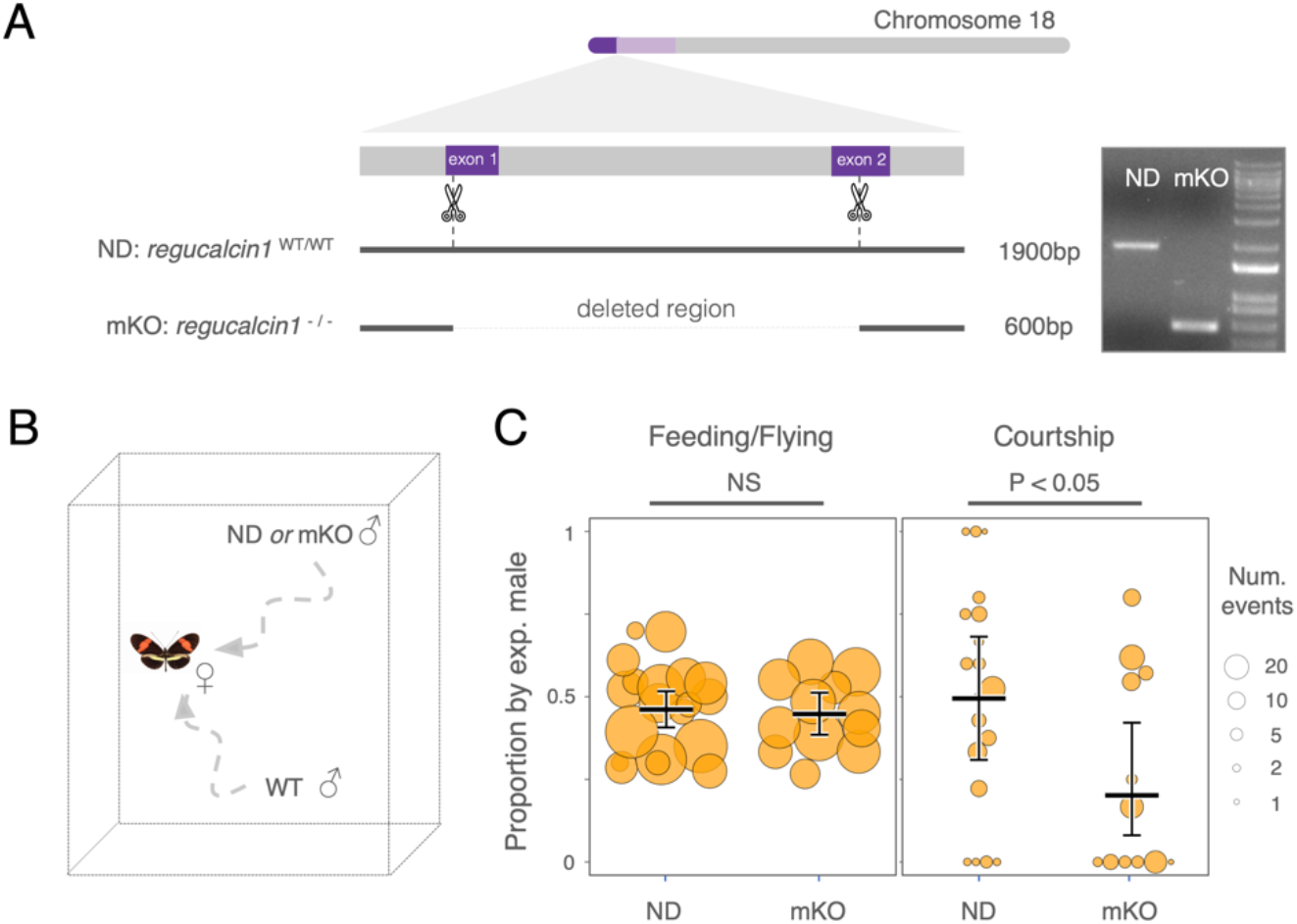
Disruption of *regucalcin1* with CRISPR/Cas9 impairs male courtship behavior. (**A**) Left: schematic representation of the *regucalcin1* locus with the target sites of the small guide RNAs and resulting CRISPR/Cas9-mediated deletion. Right: gel electrophoresis of PCR-amplified *regucalcin1* fragments from individuals without (ND) and with deletion (mKO) at *regucalcin1*. (**B**) Schematic representation of courtship trials. Experimental (*i.e.,* a mKO or ND) males that passed our ‘drop test’ were paired with a wildtype (WT) male and introduced into a cage with a wildtype virgin *H. melpomene* female. This paired design allowed us to control for both the injection procedure, as well as prevailing conditions that might potentially influence male behavior. (**C**) Proportion of time spent flying or feeding by injected but non-deletion (ND) males and *regucalcin1* mosaic knock-out (mKO) males relative to wildtype (WT) males (left panel); proportion of courtship time directed towards the same *H. melpomene* female by injected but non-deletion (ND) males (left) and *regucalcin1* mosaic knock-out (mKO) males relative to wildtype (WT) males (right panel). Point size is scaled to the number of total minutes a male flew/fed or courted during the experiments.

## Conclusions

Hybridization has been suggested to be an important source of genetic variation on which selection can act, including during behavioral evolution (*33, 34*), but direct links between specific causal genes and behavioral phenotypes are lacking. Our results strongly suggest that *Heliconius timareta* acquired a *regucalcin1* allele by hybridizing with its closely related co-mimic *H. melpomene*, increasing attraction towards red females (and presumably reproductive success). In contrast, where *H. melpomene* co-occurs with the equally closely related but differently colored *H. cydno*, *regucalcin1* contributes an important barrier to interspecific gene flow through its contribution to divergent mating preferences (*11, 35*). As such, the evolutionary impact of *regucalcin1* depends on the local mimetic landscape, emphasizing the complex role that hybridization may have on population divergence by reassembling genetic variants (*36*).

We also show that although variation in red color cue and preference map to the same genomic region, they are encoded by separate loci regulating the expression of *optix* (*15*, Fig. S10) and *regucalcin1*, respectively. By ensuring robust genetic associations between components of reproductive isolation, physical linkage is expected to facilitate speciation with gene flow, and this is likely the case for the differently colored species *H. cydno* and *H. melpomene* (*11*). However, our present results suggest these loci can also evolve independently, and evidence of a recent selective sweep in *H. timareta* at *regucalcin1*, but not *optix*, as well as distinct peaks of admixture between red-preferring species at these two genes, suggest separate introgression events. It seems likely that the acquisition of red patterns in *H. timareta* was immediately advantageous given strong selection for mimicry of local warning patterns, whereas the corresponding male preference would become advantageous only when conspecific red females had already increased in frequency.

Other prominent examples of visual preference evolution have emphasized the role of selection imposed by the broader sensory environment. In cichlid fish, for example, divergent mating preferences may have evolved as a by-product of environmental selection acting on visual pigment genes (*15, 37*). Interestingly, *H. timareta* and *H. melpomene* have evolved parallel visual preferences despite inhabiting divergent light environments (*H. timareta* is found in similar forest habitats to *H. cydno*), to which the neural and sensory systems are otherwise adapted (*38*). This suggests that visual preference evolution in *Heliconius* is not the by-product of divergent selection imposed by the broader sensory environment, but rather a consequence of direct selection to find receptive females, perhaps strengthened through reinforcement (where selection favors increased premating barriers to avoid the production of less fit hybrids)(*17, 39, 40*).

Overall, our study suggests that the evolution of *cis*-regulated differences in *regucalcin1* expression contributes to divergent mating preferences in *Heliconius*, and that hybridization can be an important source of genetic variation during behavioral evolution. The function of *regucalcin* has not been well characterized though it seems to be involved in calcium homeostasis and signaling (*41*). Our CRISPR-mediated *regucalcin1* knock-out impaired survival and flight in a few mosaic butterflies, supporting a broad role across biological processes. However, in other mosaic knock-out individuals we observed a significant reduction in mate attraction behaviors, independent of more general impairment of motor activity, implying specific effects on male mating behavior. *Regucalcin1* expression differences, sustained in adult brain tissue, likely alter how visual information is processed or integrated in the brain to determine divergent mating preferences. The challenge now is to determine the molecular and neural mechanisms through which it acts.

## Materials and Methods

### Butterfly stocks

Genetic crosses and preference trials were conducted at the Experimental Station José Celestino Mutis - Universidad del Rosario in La Vega (Colombia), between September 2019 and May 2022. Butterfly stocks for behavioral experiments were established from individuals caught around La Vega (*H. cydno cydno;* 5.0005° N, 74.3394° W) and Mocoa (*H. melpomene bellula* and *H. timareta tristero*; *1.1478° N, 76.6481° W*) in Colombia, and were maintained under common garden conditions. Larvae were reared on *Passiflora* leaves until pupation and adult butterflies were provided with ∼10% sugar solution daily and *Psiguria* flowers as a source of pollen.

### Male preference trials

We assayed preference behaviors for a total of 794 individual males across 3637 standardized choice trials (*11*). This included pure *H. melpomene bellula, H. timareta tristero, H. cydno cydno* males, as well as first generation (F1) *H. timareta tristero* x *H. cydno cydno* hybrids (obtained by crossing a *H. timareta tristero* male to a *H. cydno cydno* female) and backcross hybrids to *H. cydno cydno*. In brief, males were introduced into outdoor experimental cages (2×2×2m) with a virgin female of each type, either *H. cydno cydno* vs. *H. timareta tristero* females, or *H. cydno cydno* painted with a clear or red marker pen depending on the experiment (see below). 15-minutes trials were divided into 1-minute intervals, where courtship (sustained hovering or chasing) was scored as having occurred or not. If a male courted the same female twice during a minute interval, it was recorded only once; if courtship continued into a second minute, it was recorded twice. Whenever possible, trials were repeated 5 times for each male. From these trials we generated a data set that includes the total number of “courtship minutes” directed toward *red* and the number of “courtship minutes” toward *white* females.

### Mimicking the *H. timareta* red forewing coloration

In addition to experiments with *H. cydno cydno* and *H. timareta tristero* females, we recorded male preference phenotypes in trials with two artificially colored virgin *H. cydno cydno* females. One female had the dorsal side of the white forewing band painted with a red marker pen (R05, Copic Ciao, Tokyo, Japan), and the other with a transparent pen (Ciao 0, Copic Ciao) as control. These markers incorporate the same solvent (Copic Ciao, pers. communication). Unlike *H. cydno cydno* whose forewing band is white, *H. timareta tristero* has a red forewing band and this difference is determined by expression differences of the gene *optix*, which determines the placement of orange or red ommochrome pigments on *Heliconius* wings (*18*). Other color pattern elements also distinguish these populations, including the white hindwing margin displayed by *H. c. cydno* and a yellow hindwing bar in *H. t. tristero.* Because it is harder to match these colors across species, and because we were specifically interested in attraction to red patterns (which are the predominant difference between *H. cydno* and *H. timareta*/*H. melpomene* warning patterns across different geographical populations), we only manipulated the forewing in our experiments.

The red marker pen was chosen from several candidates (R14, R17, R27, R29, R35, R46 and RV29, Copic Ciao) to best mimic the forewing color of *H. timareta tristero* with regard to *Heliconius* color vision models. For this, we took photographs of red painted wings of *H. cydno cydno* and of *H. timareta tristero* with a Nikon Nikkor D7000 camera (Nikon, Melville NY, USA) with a visible light (380-750nm range allowed) and a UV (100–380 nm) filter in RAW format. A 40% gray standard was included in each photograph for color calibration. The visible light and UV images of each wing were combined to generate a multispectral image, using the “Image calibration and analysis toolbox” (*42*) in ImageJ (*43*). The reflectance spectra of the forewing bands were extracted from the images and converted to quantum catch models (*42*) based on photoreceptor sensitivities of *H. erato* (*44*) and relative abundance of cone receptors for species in the *melpomene/cydno* group (*44*) (*H. erato* was the only *Heliconius* species for which photoreceptor cell sensitivities had been reported at the time of this analysis). Note that *Heliconius* can discriminate in the red-range even though they have only one long-wavelength (LW) opsin with peak sensitivity at 560nm due to the presence of red-filtering pigments in some ommatidia (*45*), that shifts the peak absorbance of some cones to ∼600nm (*46*). However, this was not modeled in a first instance because the relative abundance of this cone receptor remains unknown (but see below).

We initially calculated pairwise “just noticeable differences" (JND) using a tetrachromatic (*H. erato*) color vision model with noise-limited opponent color channels, after (*47*), between the forewing band of *H. timareta tristero* and the red-painted *H. c. cydno* band using a Weber fraction of 0.05. The marker R05 had the lowest pairwise JND (0.89) and was therefore the marker we used to manipulate the forewing colors in experimental *H. cydno* females. A JND value less than 1 is considered to be generally indistinguishable by visual systems (*48*). To further corroborate that *Heliconius* males perceive the artificial and natural red patterns similarly, we acquired reflectance spectra of the artificial (red and clear) and natural (red and white) pattern elements using an Ocean Optics FLAME-T-XR1-ES spectrometer, a UV/Vis bifurcated fiber and a PX-2 Pulsed Xenon Lamp. A spectralon white standard (Ocean Optics WS-1) was used to calibrate the spectrometer. Each color pattern (*i.e.*, the forewing bar) was measured at three different locations (using an average of three scans), and the mean of the three measurements was used for further analyses. The reflectance data was analyzed through a tetrachromatic color vision model incorporating more recently published *H. melpomene* photoreceptor cell sensitivities (*49*). This differs from the model above in that we removed one UV channel and added the chromatic channel (red-shifted; λ_max_ = 590) linked to the presence of red filtering pigments (*49*) (UV-Rhodopsin1 (λ_max_ = 360 nm), blue-Rhodopsin (λ_max_ = 470nm), long wavelength-Rhodopsin without filtering pigments (λ_max_ = 570nm)). Photoreceptor cell abundances are not available for this newly classified photoreceptor type so we were unable to calculate JND values. Nevertheless, the reflectance spectra of the artificial and natural patterns overlap in shape and in tetrahedral color space when viewed under standard daylight (illum = "D65") against green foliage (bkg = "green") and with von Kris color correction (vonkries = TRUE; Figs. S1A and S1B respectively).

### Genotyping of backcross hybrids

Genotypes at the QTL peak (i.e., the region of strongest statistical association) for variation in preference behavior between *H. cydno* and *H. melpomene* on chromosome 18 (*11*) segregates with the presence of the red forewing band in our crosses due to tight linkage with the major color pattern gene *optix*. Because the presence of the red band is dominant over its absence, we were able to infer genotype at the *optix* locus by inspecting the forewing band color in backcross to *H. cydno* hybrids (*25*). Specifically, hybrid individuals with a red band are heterozygous for *H. timareta/H. cydno* alleles, and individuals lacking it are homozygous for the *H. cydno* allele. This allows a conservative test of whether this genomic region at the end of chromosome 18 influences variation in male preference based on wing pattern phenotype alone (*25*). Nevertheless, to confirm the segregation of *optix* alleles with red-color pattern in hybrid crosses and assay more specifically the genotype of hybrids at tightly linked candidate genes in the QTL peak on chromosome 18, we performed PCR amplification of a *regucalcin1* segment (found within the QTL peak). Analysis of whole genome sequence data (see below) identified indels differentiating *H. timareta* and *H. cydno* in this region, so we designed primers to encompass these putative indels at the level of *regucalcin1* (Table S2A). Genomic DNA (gDNA) was extracted from thorax tissue of our cross (*H. cydno cydno* and *H. timareta tristero*) grandparents, (*H. cydno cydno* and F1) parents and backcross hybrid progeny, using a DNAeasy Blood & Tissue kit with RNase A treatment (Qiagen, Valencia, CA, USA). Samples had previously been stored in 20 % DMSO, 0.5 M EDTA (pH 8.0) solution. We found that *H. cydno* and *H. timareta* individuals consistently differed in size of the PCR-amplified fragment, allowing us to infer genotype in the hybrid progeny. Similarly, we found indels that differentiate the two species within the QTL peak on chromosome 1 allowing us to infer genotype at this chromosomal location as well (Table S2A).

### Behavioral data analysis

We fitted generalized linear mixed models (GLMM) with binomial error structure and logit link function implemented with the R package lme4 to test for the effect of species or genotype on male preference. Specifically, we modeled the response vector of the number of “courtship minutes” toward the ‘red’ female (i.e., the *H. timareta tristero* or a red painted *H. cydno cydno* female) versus “courtship minutes” toward the ‘white’ (i.e., the *H. cydno cydno* or transparent painted *H. cydno cydno* female) and included type (i.e., species or genotype) as fixed factors. Significance was then determined by comparing models with type included as a fixed factor to models in which it was removed using likelihood ratio tests. An individual level random factor was included in all models to account for overdispersion, e.g.(*50*). Estimated marginal means and their confidence interval were extracted with the R package *emmeans*.

For our analysis testing the effect of genotype at the end of chromosome 18 on preference towards *H. timareta tristero* vs. *H. cydno cydno* females, we used the full data set of all 157 backcross males that courted *H. timareta* or *H. cydno* at least once during the trials. Genotype was initially determined from forewing color, but we updated this for 3 males of 130 males successfully genotyped at *regucalcin1* (found within the QTL peak), where we detected recombination between *regucalcin1* and *optix (i.e., optix cydno-cydno* – white forewing, QTL peak/*regucalcin1 timareta-cydno).* We note that any recombination between these loci in the individuals that we were unable to successfully genotype at *regucalcin1* will be rare (we expect just 0.62 recombination events between these two loci across the remaining 27 individuals that we could not genotype).

Although we were primarily interested in the effect of the QTL on chromosome 18, which has explicitly been shown to influence differences in visual preference between *H. cydno* and *H. melpomene* (*25*), two additional QTL have been implicated in variation in male mating preference between *H. cydno* and *H. melpomene* (*11*). The associated 1.5 lod confidence region of one of these incorporates the whole of chromosome 17, and in general is less well supported (*11, 26*). However, another behavioral QTL can be localized to a specific region of chromosome 1, for which we were able to generate genotypes (see above). To additionally include this in our analysis, we repeated our analysis of the backcross hybrids, but this time using a reduced data set including only individuals that we were able to genotype successfully at previously identified QTL on chromosome 1 and 18 (see above for details). This time the model included two fixed factors (genotype at the chromosome 18 QTL, and genotype at the chromosome 1 QTL); significance was determined by dropping each in turn and once again assessed with likelihood ratio tests. There were no quantitative differences from our previous analysis for the QTL on chromosome 18. In contrast, there was only very limited support that the QTL on chromosome 1 influences preference differences between *H. timareta* and *H. cydno* (*n* =128, 2Δln*L* = 3.79, *P* = 0.0515), and as such we did not include this QTL in subsequent analysis. Finally, in our analysis considering preference by backcross hybrids towards red and transparent colored *H. cydno* females, we again used forewing color to determine genotype at the end of chromosome 18.

### gDNA extraction and whole-genome resequencing

gDNA was extracted from thorax tissue of 4 *H. melpomene bellula* and 11 *H. t timareta tristero* individuals as well as the parents of F1 hybrids (2 *H. t timareta tristero,* 2 *H. cydno cydno,* 2 *H. melpomene rosina,* 2 *H. cydno chioneus*, see below), that were previously stored in 20 % DMSO, 0.5 M EDTA (pH 8.0) solution, using a DNAeasy Blood & Tissue kit, with RNAase treatment (Qiagen). Illumina whole-genome resequencing libraries were prepared and sequenced at Novogene (Hong Kong, China) in 125bp or 150bp paired-end mode (two different batches for *H. timareta tristero* individuals, respectively 9 and 2 samples). Previously compiled and published whole-genome resequencing data were retrieved for 5 *Heliconius numata*, 4 *H. melpomene bellula*, 10 *H. cydno chioneus*, 10 *H. cydno zelinde*, 10 *H. melpomene. rosina*, 10 *H. melpomene amaryllis* and 10 *H. timareta thelxinoe* (*19, 29, 51, 52*). Whole-genome resequencing reads were mapped to the *H. melpomene* genome version 2 (*53*) with BWA mem v.0.7.15 (*54*). Duplicate reads were marked with Picard (https://broadinstitute.github.io/picard/) and variant calling was performed with GATK v3.7 HaplotypeCaller (*55*) with default parameters except heterozygosity set to 0.02 (parameters as in (*29*), for comparable analyses). Individual genomic records were combined and jointly genotyped (GATK’s GenotypeGVCFs) for each subspecies.

### Admixture proportions, F_ST_ and d_xy_ calculation

We calculated *f*_d_ (*15*), an estimate of admixture proportion based on the ABBA-BABA test, between *H. melpomene* and *H. timareta* populations as in (*29*) and implementing scripts available at https://github.com/simonhmartin/. For this, variant sites had to be biallelic SNPs (no indels), with Quality (Q) >30 and read depth (DP) >8. In addition, variant sites were filtered out if > 30% of individuals had missing genotype calls and if > 75% of individuals had heterozygous calls. The following populations were used to estimate admixture proportions: *H. cydno chioneus* and *H. cydno zelinde* as a (combined) allopatric control population, *H. timareta tristero* and *H. melpomene bellula* (or, in a separate analysis, *H. timareta thelxinoe* and *H. m. amaryllis*) as the two sympatric species, and *H. numata bicoloratus* as the outgroup. *f*_d_ was calculated in 20kb sliding windows (step = 5kb). For *f*_d_ estimates, only sites where >60% of individuals had a genotype were considered and *f*_d_ values had to be based on >=300 ABBA-BABA informative sites per window. We also calculated sequence divergence (d_xy_) (*56*) and the fixation index (F_ST_) (*57*) in sliding 20kb windows (step = 5kb, 2000 genotyped sites required per window) with the script ‘popgenWindows.py’ available at https://github.com/simonhmartin/.

### Topology weighting

To quantify phylogenetic relationships between species in genomic intervals along the QTL region associated with visual preferences, we used *Twisst* (*30*). We used the same invariant/variant sites filtered as above (for *f*_d_ estimation), with the further requirement that at each site no more than 25% of individuals were permitted missing genotypes. Genotypes were phased and imputed using Beagle (*58*). Neighbor-joining trees (*59*) were inferred using PhyML (*60*) (substitution model = GTR), as implemented in *Twisst*. Weightings for 15 possible topologies (rooted with *H. numata* as the outgroup) were estimated for non-overlapping 50 SNPs windows.

### Selective sweeps

Variant sites were filtered for genotype quality (GQ) > 30 and read depth (DP)>10, and required to be biallelic SNPs (no indels). Furthermore, variant sites had to be called in 8 individuals out of 10 for the focal population, and in 3 individuals out of 5 for the outgroup. Sites were polarized (ancestral vs. derived) using *H. numata* as an outgroup. The background site-frequency-spectrum (SFS) was computed across the whole-genome with the exception of the Z chromosome. We used SweepFinder2 (*31*), which has been previously used to detect introgressed sweeps at color pattern loci in *Heliconius* (*61*), to estimate the composite likelihood ratio (CLR) of a sweep model compared to a neutral model (neutrality is represented by the background SFS of the genome) in 50bp steps, using both polymorphic sites and substitutions (*62*). We considered those regions with top 1% quantile CLR values as having undergone a putative selective sweep.

### Brain tissue collection, RNA extraction and sequencing

Brain (optic lobes and central brain) and eye (ommatidia) tissue were dissected out of the head capsule (as a single combined tissue) of sexually naive, 10-days old males, in cold (4 °C) 0.01M PBS. We sampled a total of 5 *H. melpomene bellula,* 5 *H. melpomene melpomene,* 5 *H. timareta tristero,* 4 *H. cydno cydno,* and *4* F1 hybrids *H. cydno cydno* x *H. timareta tristero*, which were stored in RNAlater (Thermo Fisher, Waltham, MA, USA) at 4 °C for 24 hours, and subsequently at −20 °C until RNA extraction. Previously compiled RNA-seq data for 5 *H. melpomene rosina*, 5 *H. cydno chioneus, 6* F1 hybrids *H. melpomene rosina* x *H. cydno chioneus* (generated with the same methods/in the same sequencing batch) were retrieved from (*26*). A further 5 *H. m. amaryllis* males were sampled from outbred stocks maintained at the Smithsonian insectaries in Gamboa, Panama. RNA was extracted and purified using TRIzol Reagent (Thermo Fisher) and a PureLink RNA Mini Kit with PureLink DNase digestion on column (Thermo Fisher). Illumina 150bp paired-end RNA-seq libraries were prepared and sequenced (in a single batch) at Novogene.

### Differential gene expression and exon usage

After trimming adaptor and low-quality bases from raw reads using TrimGalore v.0.4.4 (www.bioinformatics.babraham.ac.uk/projects), RNA-seq reads were mapped to the *H. melpomene* v. 2 genome (*53*)/ *H. melpomene* v. 2.5 annotation (*63*) using STAR v.2.4.2a in 2-pass mode (*64*) with default parameters (at first, see below). Only reads that mapped in ‘proper pairs’ were kept for further analysis using Samtools (*65*). For gene expression analyses, the number of reads mapping to each annotated gene was estimated with HTseq v. 0.9.1 (model = union) (*66*). For exon usage analyses, the number of reads mapping to each annotated exon was estimated using the python script “dexseq_counts.py” from the DEXSeq package (*66*). Differential gene expression analyses were conducted with DESeq2 (*67*), differential exon usage analyses with DEXSeq (*66*). Pairwise transcriptomic comparisons were conducted only between species raised in the same insectary locations (either Panama or Colombia) to avoid the confounding effect of environmentally-induced gene expression changes (Fig. S5). We considered only those genes showing a 2-fold change in expression level at adjusted (false discovery rate 5%) p-values < 0.05 (Wald test) to be differentially expressed.

An initial finding that all red-preferring subspecies showed a significantly higher expression of the last exon (5) of *regucalcin1* (HMEL013551g4) compared to white preferring species, prompted us to study whether the highly divergent sequence of red-preferring (including the *H. melpomene* reference genome) vs. white-preferring subspecies in this region might have affected this. In fact, when using more permissive parameters than the default parameters in STAR v.2.4.2a (see below), differential usage of exon 5 of *regucalcin1* disappeared in many comparisons. Given that i) with these permissive parameters there is uniform RNA-seq reads coverage of exon 5 in *H. cydno* subspecies ii) when using even more permissive parameters (parameters set 2, see below) the results remain unchanged, and that iii) when using PacBio RNA long-read data from *H. cydno* to assemble the *regucalcin1* transcript, exon 5 is included (see below), we concluded that the more permissive parameters are more appropriate, and that, the initial finding of consistent differential exon 5 usage is likely an artifact of too stringent (default) mapping parameters. We find no consistent significant changes in exon usage across all comparisons with these new parameters.

### RNA-seq mapping parameters

The default mapping parameters in STAR v.2.4.2a (*64*) were changed to more permissive ones (parameters set 1):

--outFilterMismatchNmax 15 --outFilterMismatchNoverReadLmax 0.1 --

outFilterMismatchNoverLmax 0.1 --outFilterScoreMinOverLread 0.5 --

outFilterMatchNminOverLread 0.5.

We also conducted the same analyses with yet more permissive parameters (parameters set 2):

--outFilterMismatchNmax 20 --outFilterMismatchNoverReadLmax 2 --

outFilterMismatchNoverLmax 0.2 --outFilterScoreMinOverLread 0.33 --

outFilterMatchNminOverLread 0.33.

### PacBio isoform sequencing

Brain (optic lobes and central brain) and eye (ommatidia) tissue were dissected out of the head capsule (as a single combined tissue) of sexually naive, 10-days old males, in cold (4 °C) 0.01M PBS. Tissues were stored in RNAlater (Thermo Fisher, Waltham, MA, USA) at 4 °C for 24 hours, and subsequently at −20 °C (Colombian samples) or −80 °C (Panamanian samples) until RNA extraction. RNA was extracted and purified using TRIzol Reagent and a PureLink RNA Mini Kit with PureLink DNase digestion on column from a pull of whole-brain and eye tissue of the same subspecies (4 *H. melpomene rosina*, 4 *H. timareta tristero* and 2 *H. cydno chioneus* male individuals) for a total of 3 libraries, one for each subspecies. Single molecule real-time (SMRTbell) libraries were prepared and sequenced at Novogene (Hong Kong, China), on a PacBio RSII platform (Pacific Biosciences, Menlo Park, CA, USA).

### Isoform assembly/discovery and transcript-guided annotation

Following the custom IsoSeq v3 pipeline (https://github.com/PacificBiosciences/IsoSeq/), Iso-Seq subreads from each library were used to generate circular consensus sequences (ccs), and polyA tails and artificial concatemers were removed (primers = 5’ AAGCAGTGGTATCAACGCAGAGTACATGGG, 3’ GTACTCTGCGTTGATACCACTGCTT). Bam files were transformed into fastq format using Samtools (*65*). Reads were mapped to the *H. melpomene* 2 (*53*) genome using *minimap2* (*68*) with default parameters for PacBio Iso-Seq (-ax splice:hq). Stringtie2 (*69*) was used to assemble de-novo transcripts, in order to conduct between-species comparison of isoform expression. However, coverage of Iso-Seq reads was low and the resulting transcriptome annotation sparse/incomplete not permitting inference of differential isoform expression between species.

### Allele-specific expression (ASE) analyses

8 parental individuals of the F1 hybrids *H. melpomene rosina* x *H. cydno chioneus* and F1 hybrids *H. cydno cydno* x *H. timareta tristero* (two broods for each F1 hybrid type), were genotyped using GATK v3.7 HaplotypeCaller. Individual genomic records were filtered with “hard-filters” following the GATK’s Best Practices. From these filtered variants, we extracted variant sites with opposite alleles between each parental pair with *bcftools intersect*. At the same time, we marked duplicate F1 hybrid RNA-seq reads with Picard v.1.8 (https://broadinstitute.github.io/picard/), applied the GATK’s SplitNCigarReads function and genotyped RNA-reads with HaplotypeCaller. We filtered out variant sites from F1 hybrid RNA-seq reads that had quality by depth (QD) < 2 and strand bias (FS) >30, and kept only biallelic heterozygous SNPs for further analysis (allele-informative sites should be heterozygous for the parental alleles).

Finally, we used GATK’s ASEReadCounter (without deduplicating RNA reads) to count how many RNA-reads mapped to either parental allele. We tested for differential allele specific expression for each gene with the model “∼0 + individual + allele” in DESeq2 (setting sizeFactors = 1, *i.e.,* without library size normalization between samples). By testing for ASE in F1 hybrids we can also confirm that known volumetric differences between *H. melpomene* and *H. cydno*/*H. timareta* (*38*) do not account for differences in *regucalcin1* gene expression.

### Immunocytochemistry

An affinity-purified polyclonal rabbit antibody against *regucalcin1* was developed with a *ThermoFisher* 70-days immunization protocol. Criterion to avoid cross-interaction with other epitopes was that less than 4 amino acids matched with another predicted protein from the *H. melpomene* genome assembly/annotation (Hmel2.5) (*53, 63*). The antigen target region is "EPGKFHLKKGALYRIDED". Antibodies were stored at −20°C in 50% glycerol.

Heads of insectary-reared *H. melpomene rosina* male individuals of 2-8 days of age were fixed in paraformaldehyde (PFA) 4% for 24 hours. Brains were dissected out of the head capsule in 0.02M PBS, removing the ommatidial, retinal and laminal tissue, and then embedded in 4% agar and sliced at 250nm with a LeicaVT1200S vibratome. As such, because our samples did not include ommatidial and laminal tissue, we cannot rule out expression in these more peripheral stages of visual processing. Brain sections were washed with blocking solution (BS: 1,5% Triton X-100 ; 0,1% Saponin; 1% bovine serum albumin) 3 times for 30 minutes at room temperature and then incubated at 4°C for 2 days with 1:100 rabbit *regucalcin1* primary antibody (we combined an equal amount of two immunized rabbits sera, with 1.24 and 2.5 mg/ml concentration respectively before glycerol dilution) and 1:30 mouse *synapsin* (anti SYNORF1, Developmental Studies Hybridoma Bank, University of Iowa, Iowa City, IA,RRID: AB_528479) in BS solution. Samples were washed 3×30min in BS at room temperature, and then incubated for 1 day at 4°C with Alexa 647 anti-rabbit (Dianova, 711-606-152, 1:300), Cy3 anti-mouse (Dianova 715-166-151, 1:400), and Neuro Trace blue (Mol probes invitrogen, 1:300) in BS. Finally, samples were washed 3×30min in PBS, and then mounted in Vectashield medium. Although this data does not pinpoint the site of action, they confirm that regulatory changes of *regucalcin1* could influence preference by affecting processing at multiple sites along the visual pathways.

### Confocal imaging and image analysis

Brains were imaged with a Stellaris 5 confocal microscope (Leica) equipped with a white light laser and a 405nm laser, and a HC PL APO CS2 40x /1.10 water immersion objective and with the tile scanning function. Excitation wavelengths and emission filters were 405 nm and 476-549 nm for neurotrace blue, 554 nm and 559-658 nm for Cy3, and 653nm and 658-750 nm for Alexa 647. Images were acquired with a pixel size of 0.142 μm and a pinhole aperture of 1 Airy unit. Confocal images were analyzed on ImageJ (https://imagej.nih.gov/ij/). A median filter was applied and signal intensity adjusted on whole images for each wavelength.

### CRISPR/Cas9-mediated mutagenesis of r*egucalcin1*

*Heliconius melpomene rosina* pupae were obtained from a commercial supplier (https://www.butterflyfarm.co.cr) and used to establish a stock in an external greenhouse at LMU Munich. We used *GeneiousPrime* v2021.1 to design 4 guide RNAs corresponding to N_20_NGG (on either strand), targeting exon1 and exon2 of *regucalcin1* (Table S2B), considering the gRNA efficiency scores predicted from (*70*), favoring GC-rich regions close to the PAM (NGG) sequence, and avoiding polymorphic sites in our butterfly stock. N_20_NGG sequences were screened for off-targets in the *H. melpomene* 2.5 genome with the BLAST function of Lepbase v4. Only guide RNAs that had unique seed regions 12bp upstream of the PAM were considered further to avoid off-targets.

Synthetic sgRNAs were ordered from *Synthego* (Redwood City, CA, US) and resuspended in TE (0.1mM EDTA, pH 8.0) buffer (Sigma Aldrich, St. Louis, MO, US). Cas9 protein (CP01, PNAbio) was reconstituted in nuclease-free water and 5% Phenol Red Solution (Sigma Aldrich), following the guidelines in (*71*). A mix of 4 gRNAs and later 2gRNAs and Cas9 protein (250:500ng/µl) was injected in eggs between 1 and 4.3 hours after laying, using a Femto Jet (Eppendorf, Hamburg, Germany).

To genotype mosaic generation zero (G0) individuals, we extracted gDNA from two caterpillar spikes at 4^th^/5^th^ instar by squishing the spikes with a filter tip in 9 µl NaOH solution (50mM), incubating at 95°C for 15 minutes, cooling the reaction on ice for 2 minutes and adding 1 µl of Tris-HCl (1M) (Nicolas Gompel and Luca Livraghi pers. comm., modified from (*72*)). We then PCR-amplified a region of *regucalcin1* (Table S2), to screen for CRISPR/Cas9 mediated deletions as a result of non-homologous end-joining following multiple double-strand breaks predicted to result in a ∼600bp DNA fragment (with deletion) instead of ∼1900bp (no deletion).

We purified DNA from gel bands of the allele carrying the predicted deletion with a MinElute Gel Extraction Kit (QIAGEN) and ExoSap (Thermo Fisher) and Sanger-sequenced with a BigDye v1.1 kit (Thermo Fisher) with the Genomics Service Unit of LMU Munich to find that the same 2 gRNAs (Table S2) consistently mediated the introduction of a deletion and were therefore used for generating *regucalcin1* mKO butterflies in all experiments (survival/efficiency statistics for CRISPR experiments in Table S1). Although most CRISPR-mediated deletions were of the expected size (1300bp deletion), in a few mKO individuals the deletion varied in size (ranging from ∼400bp to ∼1500bp), probably due to variation in the DNA repair process. Nevertheless, we found that the boundaries of these deletions always coincided with either one of the two sgRNA target sites, likely generating similarly non-functional alleles. Individuals that were screened as mKO at the 4th/5th instar were subsequently confirmed as mKO by PCR on DNA extracted from adult brain, thorax or abdomen tissue with a DNAeasy Blood & Tissue kit.

We extracted gDNA from at least two tissues among brain, thorax and abdomen from 40 individuals without deletion (ND), and sequenced their *regucalcin1* protein-coding region to screen for small frame-shift mutations/deletions following double-strand breaks at only one of the CRISPR target sites, which would not be detected by our PCR-fragment size screen. We found that only 1/40 individuals (2.5%) showed evidence of a CRISPR-mediated mutation at only one of the target sites not resulting in a large deletion. Thus, ND mKO individuals with small frame-shift mutations are rare and might have only marginally impacted the results (i.e., considered as ND instead of mKO). On the other hand, mKO individuals had a substantial percentage of cells carrying the deletion in their brain tissue (Fig. S8).

### Drop test

To assay basic locomotor (flying) function of *regucalcin1* mKO butterflies, we conducted a ‘drop test’ with mKO, ND or WT butterflies one day post-eclosion during the butterfly’s active hours (between 10:20 and 17:30). Each butterfly was held by the forewings 1.5 m above the ground at the center of a 2×2×2m cage and then released. This procedure was repeated 5 times for each butterfly, and individuals were considered to have ‘failed’ the test if they dropped directly on the ground (instead of flying) for all 5 trials (examples negative and positive responses, Video S1-S2). With the exception of three individuals (one mKO, one ND and one WT), all butterflies either dropped to the ground on all 5 trials, or flew on all five trials.

### Optomotor assay

To determine whether *regucalcin1* mKO butterflies show a visual (optomotor) response, *i.e.,* an innate orienting response evoked by wide-field visual motion, we placed mKO, ND or WT butterflies >4 hours post-eclosion at the center of an experimental arena of 16 cm radius surrounded by a visual stimulus of alternating black and white stripes (*73, 74*). We used a visual stimulus with spatial frequency value (cycles-per-degree) of 0.2 cycles-per-degree (cpd). The width (in millimeters) of one cycle (a set of alternating black and white stripes) was calculated as *cycle width = [(C/360) / a]*, where ‘*C*’ is the circumference of the experimental arena (mm) and ‘*a*’ is the visual acuity (cpd). Butterflies were restrained in a clear PLEXIGLAS® cylinder and all assays were conducted at room temperature under illumination from an overhead LED lamp, and recorded with a GoPro camera (GoPro, San Mateo, CA, US) placed above the device. Butterflies were tested once they stopped crawling on the cylinder, which was followed by 6 rotations of the stimulus (alternating between clockwise and counterclockwise), each lasting 10 seconds, and running at a speed of three rotations per minute (3 rpm). A positive response was scored if the butterfly changed the orientation of its head/antenna in the direction of the moving stimulus and then re-oriented itself in the opposite direction when the direction of rotation was reversed, across the whole 1-minute trial (see Video S3 for an example).

### Courtship assay

mKO, ND and WT *H. melpomene* males were maintained together in a 2×2×2m cage in a greenhouse in Munich. As in our experiments in the tropics, butterflies were provided with *Lantana* and *Psiguria* flowers, as well as a sugar water supplement daily. All courtship trials were conducted between 11:00 and 1700. We paired either an experimental mKO or ND >5days post explosion male with a WT male (matched for age, but otherwise chosen at random). This paired design allowed us to control for both the injection procedure, as well as prevailing conditions that might potentially influence male behavior. Individuals that failed the drop test were excluded from courtship assays (as none survived five days post-eclosion). A WT virgin *H. melpomene* female (1-5 days post eclosion) was then introduced into the cage. As for our behavioral experiments in Colombia, 15-minute trials were divided into 1-minute intervals, and during each minute both the experimental and WT male were scored for three behaviors flying, feeding, and courting (sustained hovering over or chasing the female for >3 seconds). The experimental cage was shaken every 5 minutes to stimulate butterfly activity. In the minute-interval following cage shaking, a flying occurrence was recorded only if it lasted for 10 uninterrupted seconds, or occurred after a butterfly had momentarily landed/stopped flying. The trials were stopped immediately if mating occurred (and the butterflies were gently separated). Trials were repeated up to 5 times for each experimental male (median=3). To further avoid biasing our results, we excluded from trials a single mKO male that did not fly, court or feed during all 4 trials in which it was tested (though this more conservative approach does not qualitatively affect our results).

As with data from our behavioral trials in Colombia, we tested for differences in relative courtship activity between mKO and ND males using generalized linear mixed models (GLMMs) with binomial error structure and logit link function (implemented with the R package lme4). This time the proportion of minutes courting females by experimental (i.e. mKO or ND) vs WT males was the dependent variable and the experimental male type (mKO or ND) was set as a fixed explanatory factor. We tested significance by comparing this model to a null model, excluding experimental type as an explanatory variable, with a likelihood ratio test. Once again, experimental male ID was included as an individual level random factor in all models to account for overdispersion. To determine whether mKO and ND males differ in more general motor activities, we repeated these analyses, but this time with the proportion of minutes spent flying or feeding by experimental versus WT males. Estimated marginal means and confidence intervals were extracted using *emmeans*.

### Patternize analysis

To determine whether *regucalcin1* mKO affects wing color patterns in *H. melpomene rosina*, we quantified and compared color patterns of mKO and ND butterflies using *patternize* (*75*). Wing pictures were taken in RAW format with a Fujifilm X-T3 camera with a Fujifilm 35mm F1.4 R lens, using a white-diffusion sheet (Lee filters 252) to homogenize lighting from two overhead LED lamps. The white balance of each image was then adjusted with the *Curves* feature (constant settings) in Adobe Photoshop CC 2019 (Adobe, CA, USA), to mask either one of the butterfly forewings (marked with a marker pen to keep track of individual butterflies ID) and to remove the background. To align wing images, we positioned 18 and 16 landmarks respectively (as suggested in the *patternize* package) at vein intersections on the forewings and hindwings for each sample (Fig. S10A). A *thin plate spline* transformation was then used to align landmarks to a common (arbitrarily chosen) reference sample for each of 4 groups (mKO males, mKO females, ND males, ND females) and each 4 patterns (forewing dorsal, forewing ventral, hindwing dorsal, hindwing ventral) independently. To compare pattern size and shape among samples, the red, green, and blue (RGB) values were extracted for each pattern of each group separately using *patternize*, with color threshold “colOffset”, and the relative size of the pattern was calculated as the proportion of the pattern area over the total wing area (of the same wing) using the *patArea* function in *patternize*. Differences in color pattern among groups were calculated by subtracting the pattern frequencies with the “sumRaster” function in *patternize*. The resulting rasters were analyzed using Gower’s dissimilarity measure (*76*) in the *R* package *StatMatch,* as commonly used for *patternize* data (for example in reef fishes, (*77*)), to determine statistically significant differences in pattern spatial distribution among groups (Fig. S10).

## Data and analysis scripts

All raw data and analysis scripts are available at: https://github.com/SpeciationBehaviour/Adaptive_introgression_of_a_visual_preference_gene. Genome re-sequencing and RNA-seq data will be published in the European Nucleotide Archive (ENA) upon acceptance for publication.

## Acknowledgements

We dedicate this paper to the memory of our friend and colleague Alexander Hausmann. We are grateful to Bianca Hoelldobler, Isabel Leon, Francesco Rossi, Rebecca Stephens, Sophie Smith, José Borrero, Marilia Freire, Alberto Comin, Christina Burrows, Michaela Bauer and Christine Rottenberger for technical and rearing assistance, and Mathieu Choteau for help with fieldwork. We thank Francesco Cicconardi for sharing a pipeline for Iso-Seq analysis and Simon Martin for sharing vcf files. We thank Steven van Belleghem, Philipp Brand, Max Farnworth, Nicolas Gompel, Joseph Hanly, Luca Livraghi, Stephen Montgomery, Ricardo Pereira, Jochen Wolf and Vera Warmuth for valuable input on methods and the manuscript. We are grateful to *Autoridad Nacional de Licencias Ambientales*, Colombia for permission to collect butterflies.

## Author contributions

R.M.M. conceived the project; M.R., A.E.H and R.M.M designed the experiments; A.E.H., D.L., L.M. G.R. and R.M.M oversaw the genetic crosses and collected the behavioral data; C-Y.K. and D.S.W. did the color vision analysis; M.L. arranged the collection of butterflies in Colombia; M.R., A.E.H and R.M.M analyzed the behavioral data; M.R. and S.M. analyzed the population genomic data; M.R. collected tissue and analyzed the transcriptomic data; M.R. and M.M. performed the selective sweep analysis; M.R. and P.A. performed brain immunostainings; P.A. confocal imaging and image analysis; M.R. performed the CRISPR-Cas9 and associated experiments; D.S.W. developed the optomotor assay; A.M performed the patternize analysis; W.O.M., C. P-D, C.S. and R.M.M. provided materials as well as supervision and insights for data collection and analysis. M.R. and R.M.M. wrote the paper with input from all authors. (**A**) Manipulation of *H. cydno* female forewing color with a red marker pen (photo credit: Tal Kleinehause Gedalyahou). (**B**) Reflectance spectra of the natural red and red-painted forewing bars, as well as of the white and clear-painted (transparent marker) forewing bars averaged across 4 *Heliconius timareta tristero,* 4 painted *H. cydno cydno*, 9 *H. c. cydno and 4* painted *H. c. cydno samples* respectively. Shaded regions represent ±1 standard error. (**C**) Tetrahedral color space, *i.e.*, predicted stimulation of different photoreceptor cell types, for the different forewing reflectances, using a tetrachromatic model with *H. melpomene* photoreceptor cell sensitivities (*49*). Corners indicate photoreceptor cell-type maximum sensitivities: UV-Rhodopsin1 (360 nm), blue-Rhodopsin (470nm), long wavelength-Rhodopsin without (570nm) and with red filtering pigments (+R) (590nm). Solid circles indicate unmanipulated forewings (n=5), open circles indicate painted forewings (n=5), and the solid square indicates the achromatic point of equal stimulation for all photoreceptors. (**D**) Proportion of courtship time directed towards red painted *H. c. cydno* females relative to white (transparently painted) *H. cydno* females, by *H. timareta* and *H. cydno* males, and **(E)** by F1 hybrids and backcross to *H. cydno* hybrid males. Orange points represent individuals that are heterozygous (*i.e., H. cyd*/*H. tim.*) and blue points represent individuals that are homozygous for (*i.e., H. cyd.*/*H. cyd.*) *H. cydno* alleles at the *optix* locus on chromosome 18 (and tightly linked regions, including the QTL peak). Dot size is scaled to the number of total minutes a male responded to either female type. Estimated marginal means and their 95% confidence intervals are displayed with black bars.

---

**Fig. S1.**
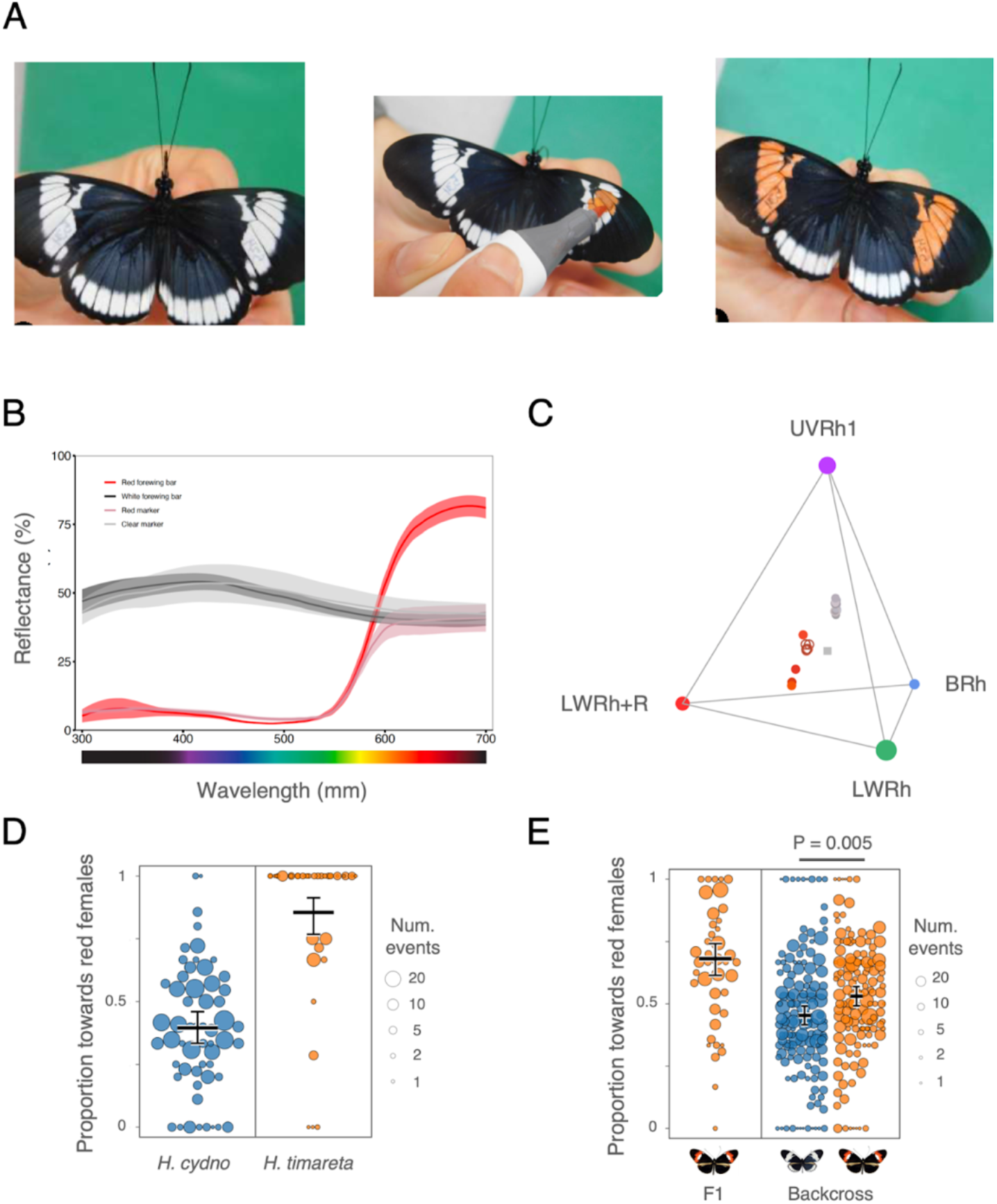
Species mating preferences and the behavioral QTL on chromosome 18 are visually guided.

**Fig. S2.**
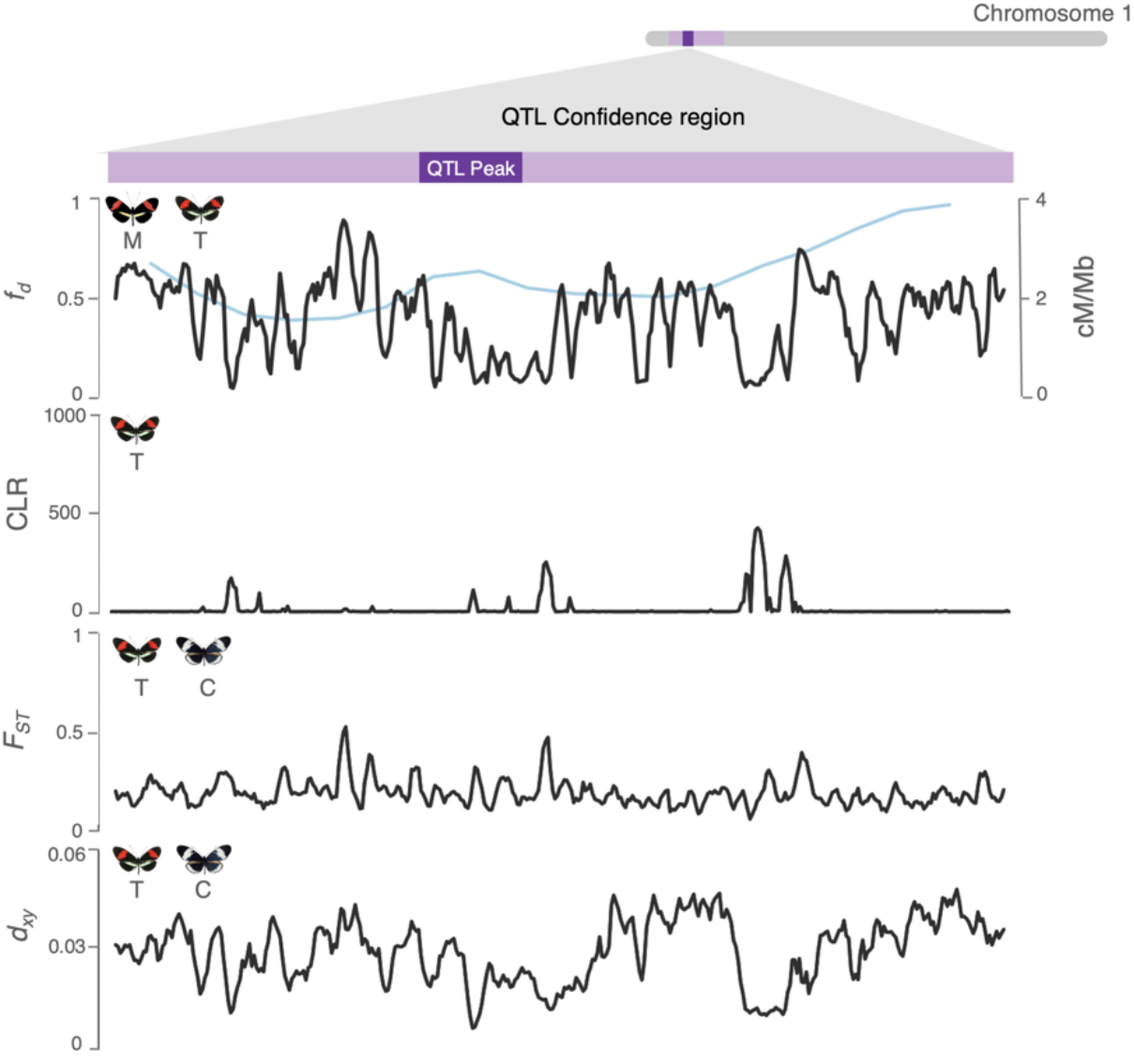
Genomic signatures of adaptive introgression and divergence at the behavioral QTL on chromosome 1. Top panel: Admixture proportion values between *H. melpomene a*nd *H. timareta* at the behavioral QTL region on chromosome 1. Recombination rates (as estimated in (*78*)) overlayed in blue. Second panel: composite likelihood ratio (CLR) of a selective sweep in *H. timareta.* Third and fourth panels display fixation index (F_ST_) and d_xy_, between *H. timareta and H. cydno*.

**Fig. S3.**
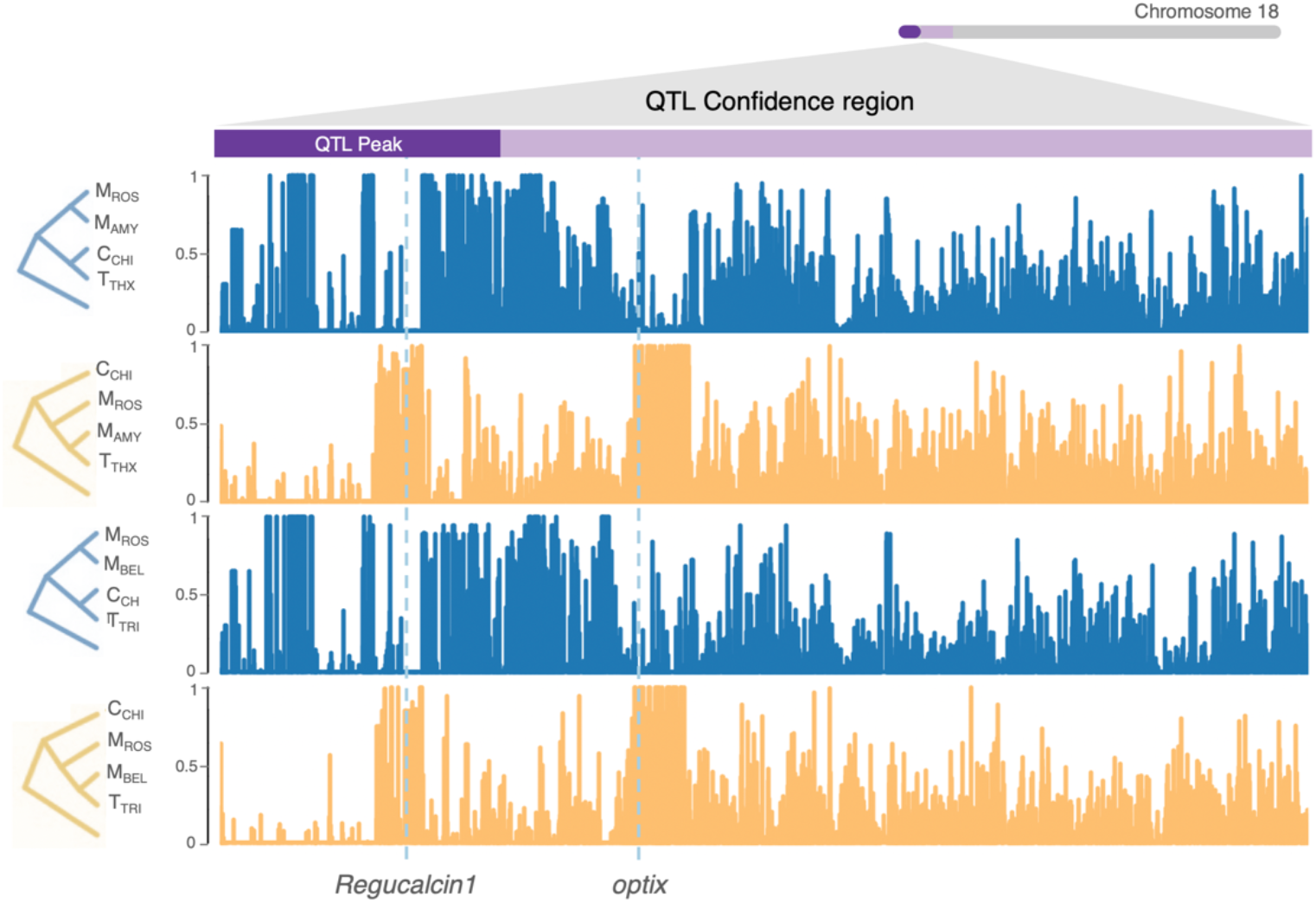
Sharing of alleles between different populations of red-preferring species at *regucalcin* and *optix*. Topology weightings, *i.e.,* proportion of a particular phylogenetic tree over all possible rooted trees, along the behavioral QTL region on chromosome 18 (*x*-axis represent physical position). The “species” tree (expected species relationships: *H. timareta* more closely related to *H. cydno* than *H. melpomene*) is represented in blue, the “introgression” tree (where *H. timareta* clusters with its sympatric *H. melpomene* co-mimic) in orange. Top two panels: focal populations of *H. timareta* and *H. melpomene* from Peru (*H. m. amaryllis* and *H. t. thelxinoe*). Bottow panels: focal populations from Colombia (*H. m. bellula* and *H. t. tristero*). *H. numata* was used as outgroup. Gene coordinates of *regucalcin1* (candidate behavioral gene) and *optix* (color pattern gene) are highlighted by vertical light blue dotted lines.

**Fig. S4.**
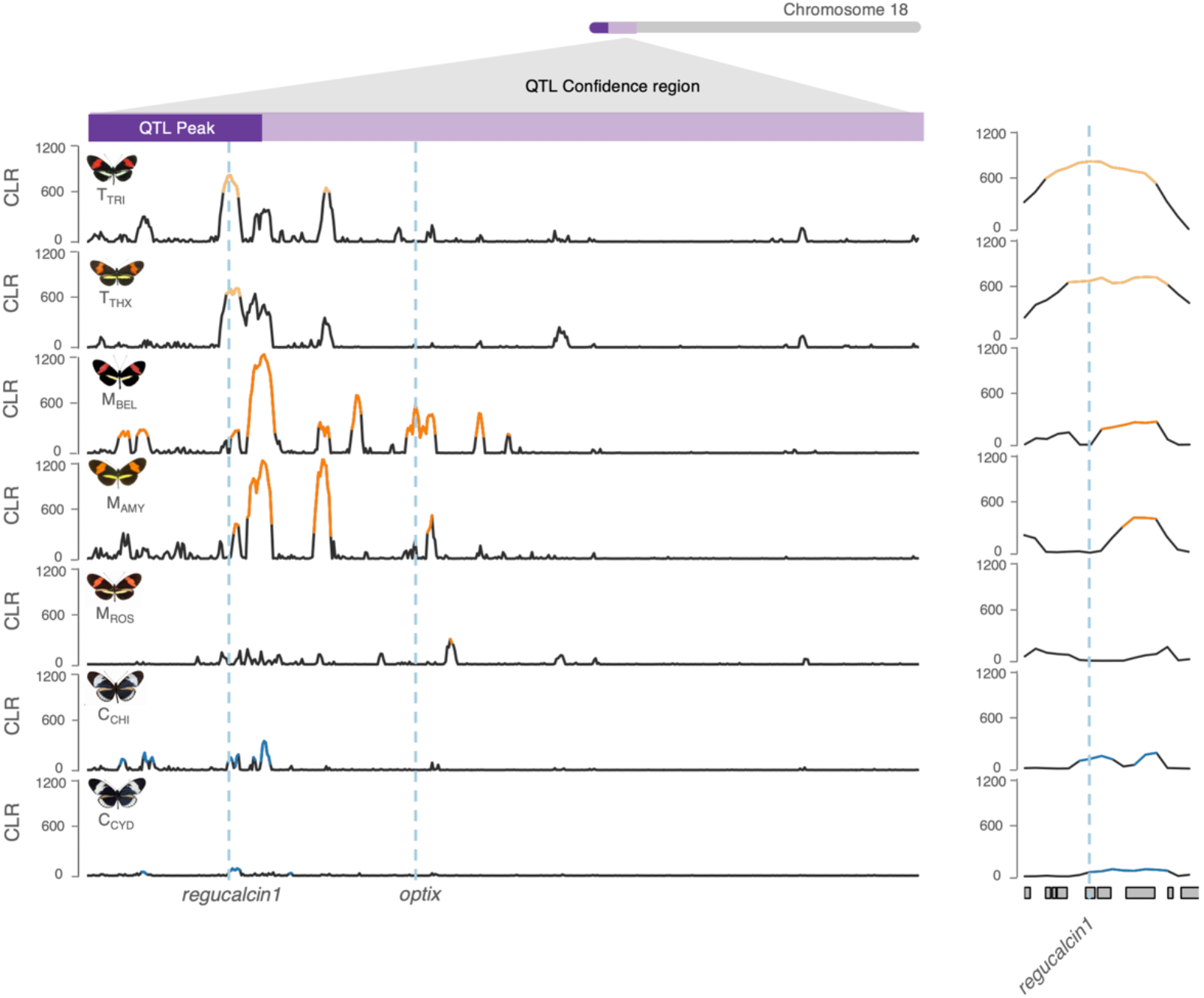
Evidence for a selective sweep at the *regucalcin* locus across different *Heliconius* populations. Composite likelihood ratio (CLR) of a selective sweep in different *Heliconius* populations across the QTL region on chromosome 18. Top 1% quantile values are highlighted with colors. Note that i) the highest support for a selective sweep in *H. melpomene* populations is centered at ∼100 kb from the *regucalcin* locus and likely represent a more recent selective sweep at a locus other than *regucalcin* ii) the considerably lower absolute CLR score in *H. cydno* populations compared to *H. timareta* populations at *regucalcin* could represent the effect of background selection (removal of deleterious variants), remnants of an old selective sweep or noise instead of positive selection. M_AMY_ = *H. melpomene amaryllis* (Peru), M_BEL_ = *H. melpomene bellula* (Colombia), M_ROS_ = *H. melpomene rosina* (Panama), M_MEL_ = *H. melpomene melpomene* (Colombia), T_TRI_ = *H. timareta tristero* (Colombia), T_THX_ = *H. timareta thelxinoe* (Peru), C_CHI_ = *H. cydno chioneus* (Panama), C_CYD_ = *H. cydno cydno* (Colombia).

**Fig. S5.**
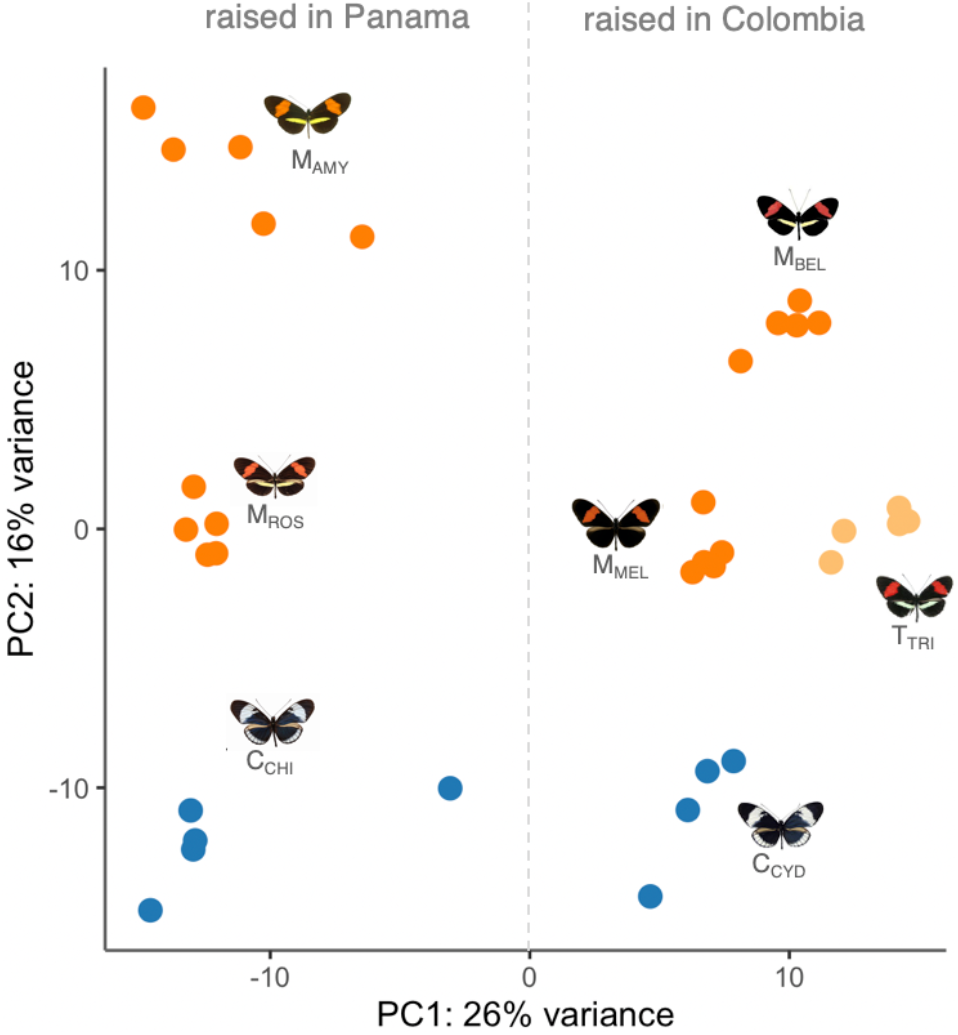
Brain and eye transcriptomic profiles cluster by rearing environment and species. Principal component analysis (PCA) of gene expression levels for the 500 genes with most variable expression level across brain tissue samples from different species. Samples are color-coded by species. A vertical dotted line has been drawn to indicate the division (PC1) between individuals that were raised in Panama (*H. c. chioneus* (C_CHI_) and *H. m. rosina* (M_ROS_) as previously described (*26*)) and in Colombia. Interestingly, *H. timareta* clusters more closely to *H. melpomene* (by visual preference phenotype) than to *H. cydno* (by phylogeny), suggesting broad convergence in neuro-transcriptomic profiles between sympatric, hybridizing populations of *H. melpomene* and *H. timareta*, raised in common garden conditions. M_AMY_ = *H. melpomene amaryllis* (raised in Panama), M_BEL_ = *H. melpomene bellula* (raised in Colombia), M_ROS_ = *H. melpomene rosina* (raised in Panama), M = *H. melpomene melpomene* (raised in Colombia), T_TRI_ = *H. timareta tristero* (raised in Colombia), C_CHI_ = *H. cydno chioneus* (raised in Panama), C_CYD_ = *H. cydno cydno* (raised in Colombia).

**Fig. S6.**
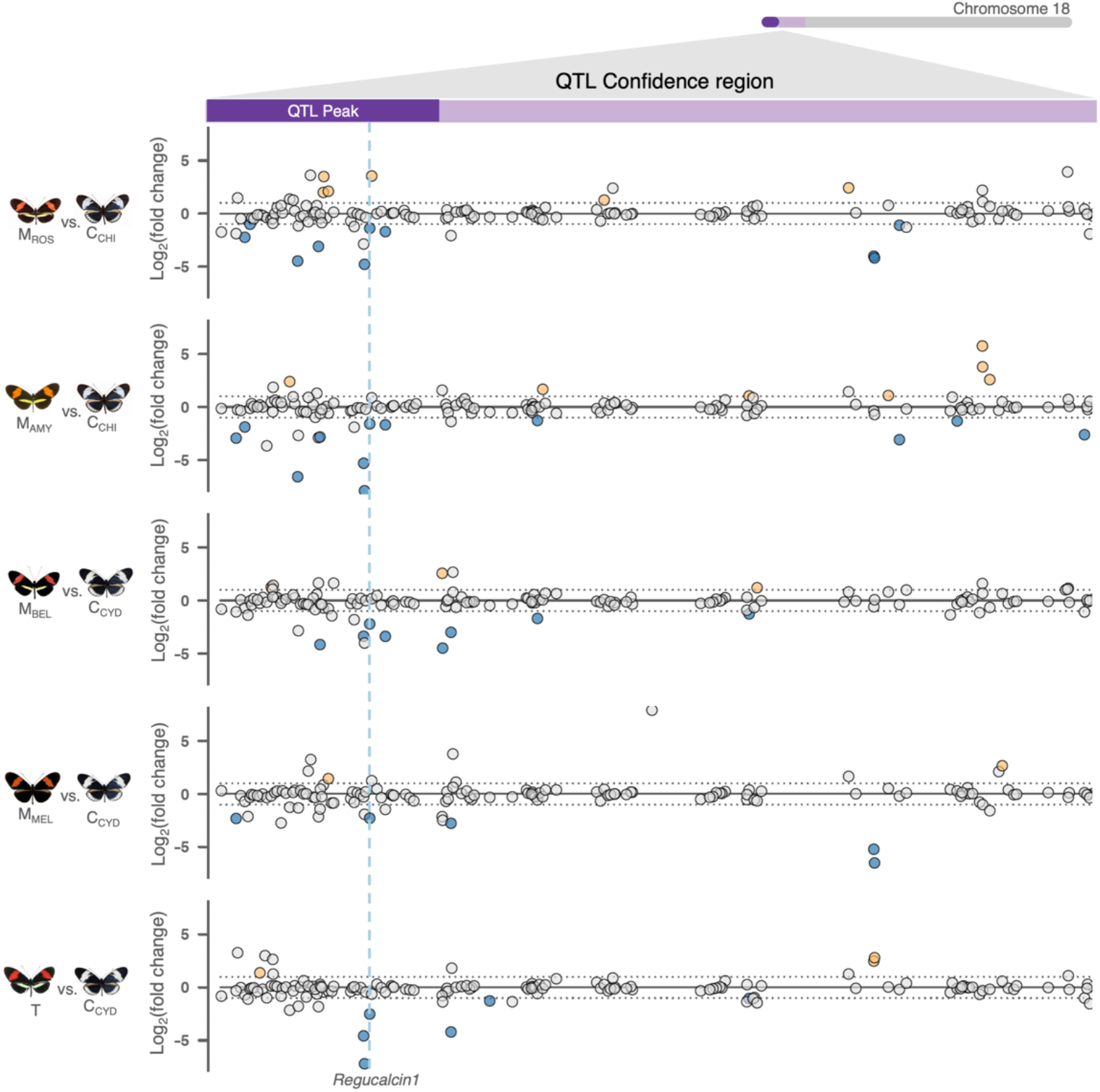
Differential expression across populations at the preference QTL region on chromosome 18. Points correspond to individual genes and the *y*-axis indicates the *log_2_* (fold-change) for each “red-preferring” vs “white preferring” subspecies comparison. The QTL peak, and the rest of the QTL confidence interval on chromosome 18 are shown on top in dark and light purple respectively (*x*-axis represents physical position). The two horizontal dashed lines (at *y*-values of 1 and −1) indicate a 2-fold change in expression. Genes showing a significant 2-fold+ change in expression level between groups are highlighted in orange and blue, where orange indicate a 2-fold higher expression level in *H. melpomene* subspecies or *H. timareta,* whereas blue a 2-fold higher expression level in *H. cydno*. A vertical dashed blue line highlights the only gene that is differentially expressed between all comparisons*: regucalcin1* (higher expression level in *H. cydno* populations). M_AMY_ = *H. melpomene amaryllis* (raised in Panama), M_BEL_ = *H. melpomene bellula* (raised in Colombia), M_ROS_ = *H. melpomene rosina* (raised in Panama), M = *H. melpomene melpomene* (raised in Colombia), T_TRI_ = *H. timareta tristero* (raised in Colombia), C_CHI_ = *H. cydno chioneus* (raised in Panama), C_CYD_ = *H. cydno cydno* (raised in Colombia).

**Fig. S7.**
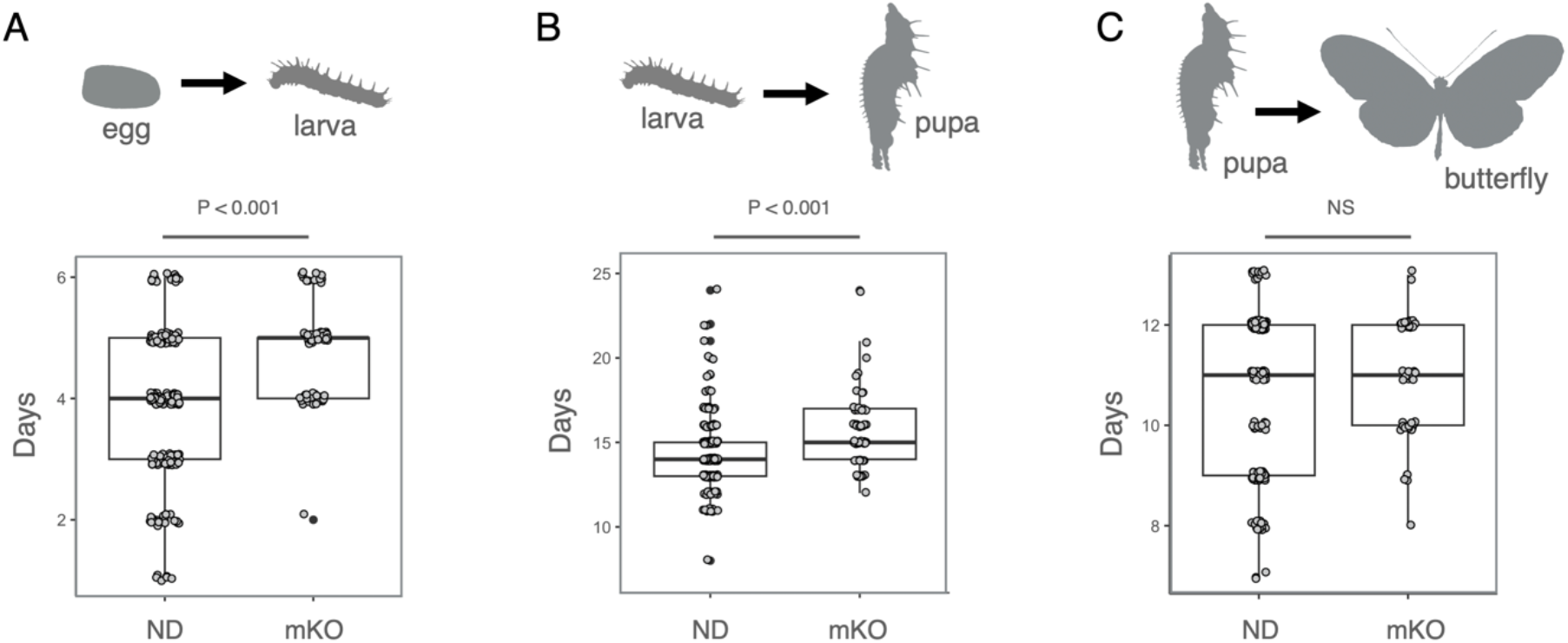
CRISPR/Cas9 mediated knock-out of *regucalcin1* delays development in its early stages. Days it took to develop **(A)** from egg to larva (unpaired *t*-test: P <0.001) **(B)** from larva to pupa (unpaired *t*-test: P <0.001) and **C)** from pupa to imago (adult) (unpaired *t*-test: P > 0.05) for individuals without (ND) and with deletion (mKO) at the *regucalcin1* locus.

**Fig. S8.**
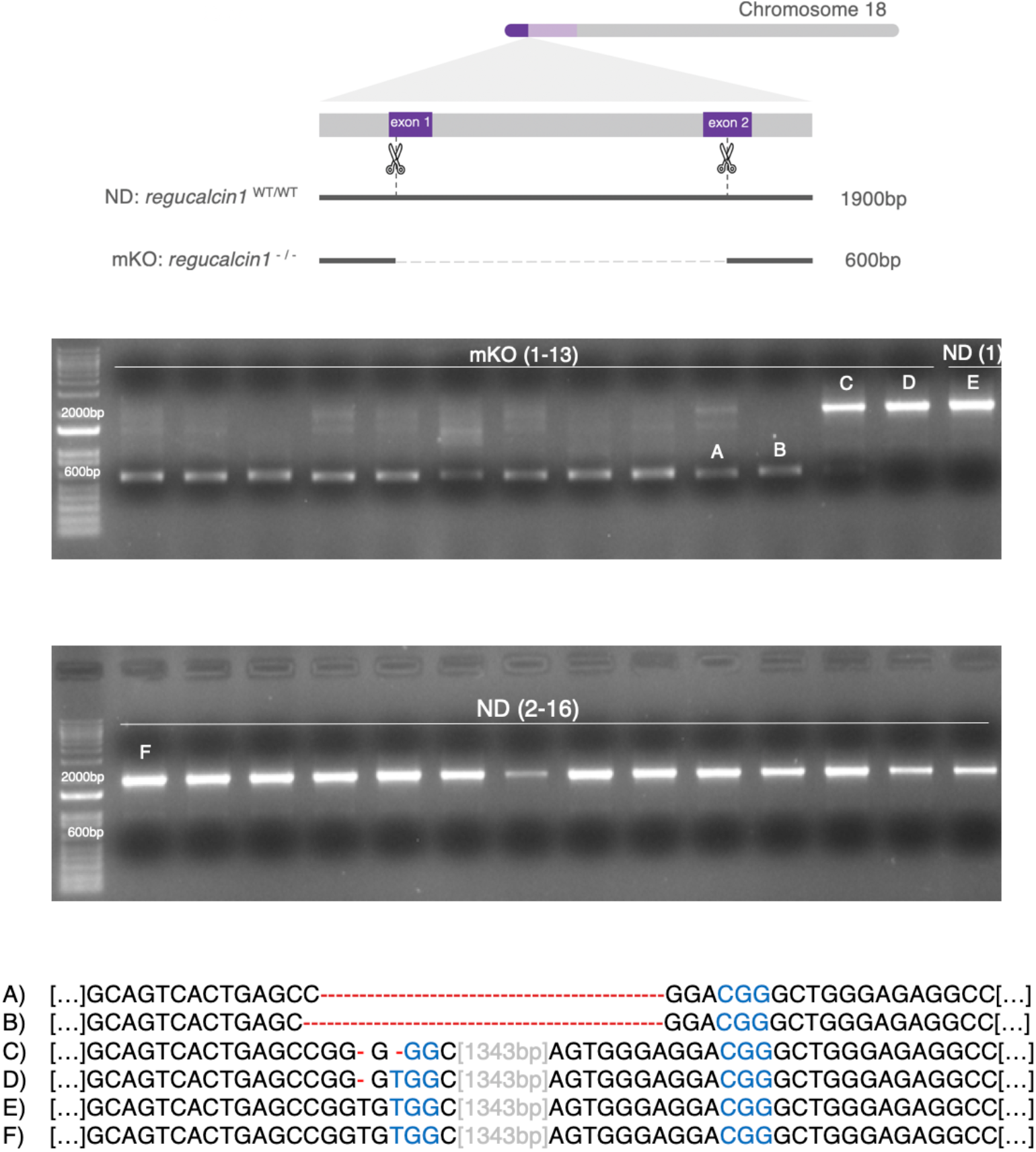
A high percentage of cells show *regucalcin1* knock-out in G0 mosaic individuals. On top, schematic representation of the *regucalcin1* locus with the target sites of the small guide RNAs and resulting CRISPR/Cas9-mediated deletion. Below, gel electrophoresis of PCR products of the *regucalcin1* locus from DNA extracted from whole brain tissue of mKO and ND males that were tested in courtship assays (note that 1 ND male sample was not included for space constraints on the gel, and that DNA extraction could not be carried out for 3 ND individuals, whose bodies could not be recovered). Below, examples of nucleotide sequences for alleles carrying and not carrying the deletion (as inferred with Sanger-sequencing of DNA purified from the respective gel bands). Note that for sample mKO 12 (C) there is also a small percentage of cells with deletion, whereas for sample mKO 13 (D) only a single-nucleotide frame-shift mutation.

**Fig. S9.**
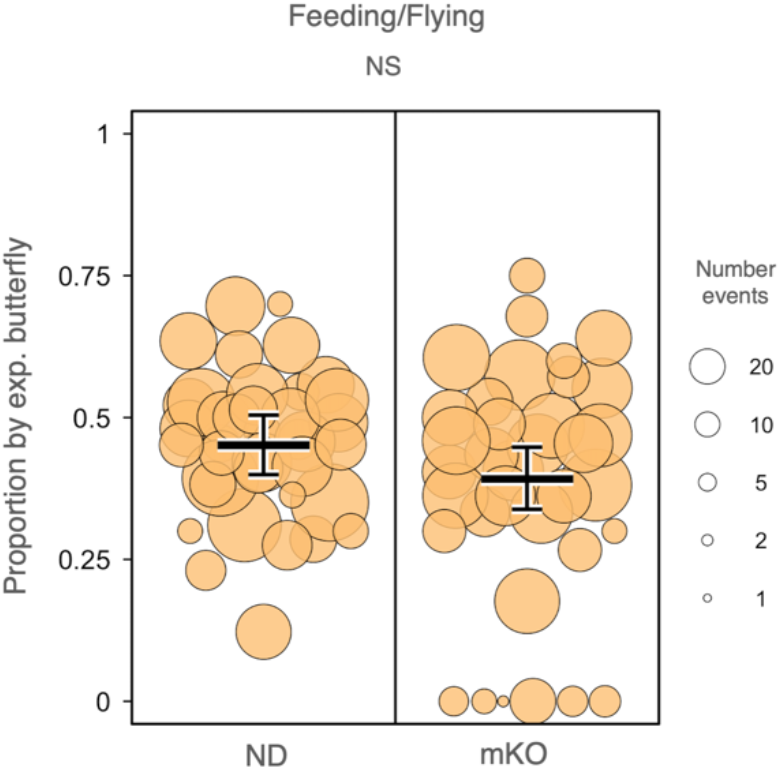
No significant change in flying and feeding behaviors caused by *regucalcin1* knock-out. Proportion of time spent flying and/or feeding by ND individuals (left) and *regucalcin1* mKO individuals (right) relative to wild-type butterflies (female and male individual tested, females were tested as of 1-day of age). These include four females that did not pass the drop test and (two additional males that) did not show any flying or feeding activity (values = 0). Dot size is scaled to the number of total minutes individuals flew and/or fed during the experiments.

**Fig. S10.**
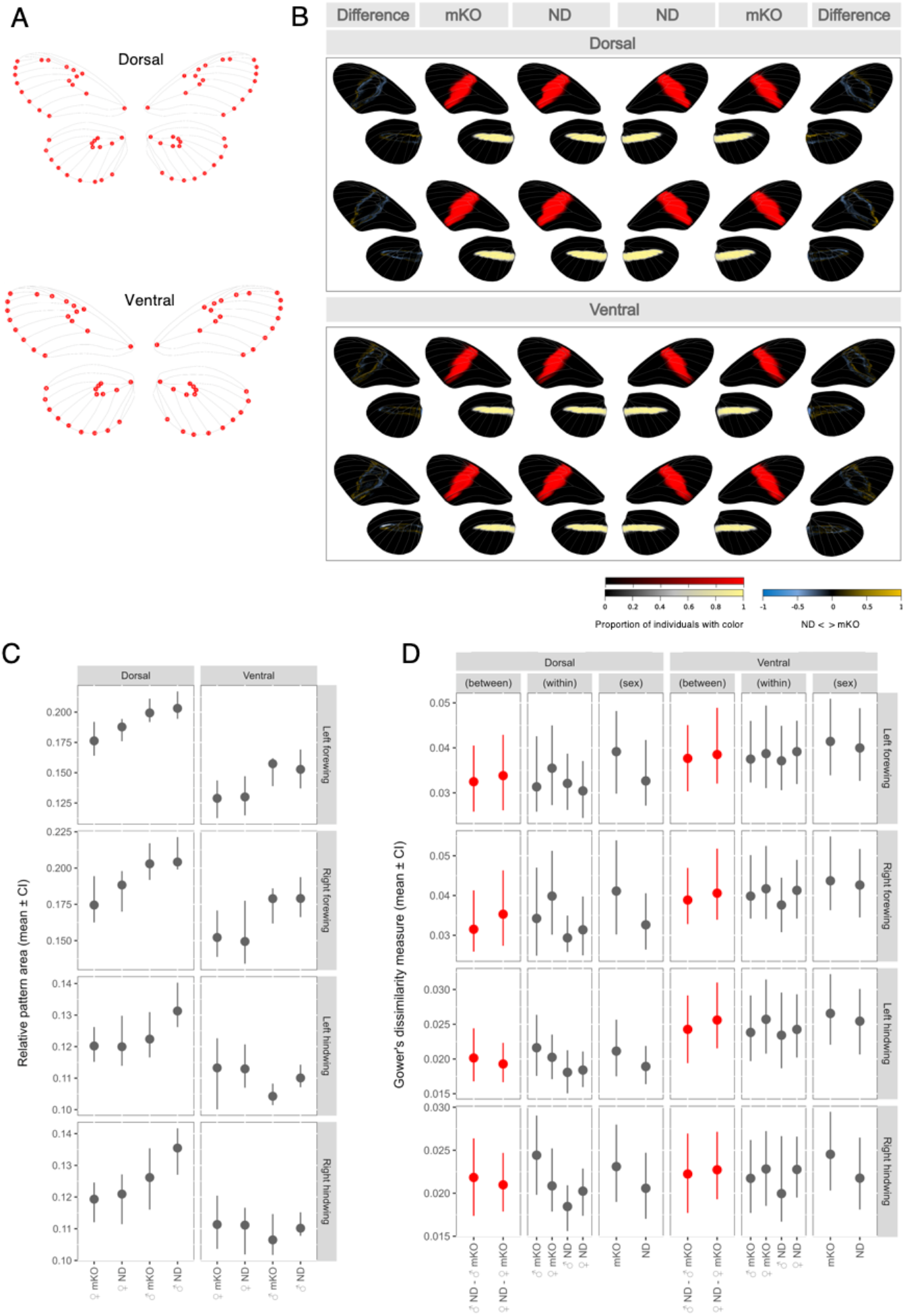
No evidence for an effect on color pattern in *regucalcin1* mKO individuals. (**A**) Landmarks placed at the intersection of the forewing and hindwing veins for dorsal and ventral wing sides. (**B**) Average color patterns (central columns) and differences in color pattern (leftmost and rightmost columns) between *H. melpomene rosina* mKO and ND (*i.e.* with and without deletion at *regucalcin1*) individuals, analyzed separately by sex, forewing (FB) and hindwing band (HB), and dorsal and ventral sides (sample sizes: 26 mKO females, 20 mKO males, 23 ND females and 19 ND males). Yellow indicates higher presence of FB/HB in mKO butterflies and blue indicates higher presence of FB/HB in ND butterflies. (**C**) Average pattern area of FB and HB for each group, with 95% confidence intervals. (**D**) Mean Gower’s dissimilarity measure of FB and HB between-group, within-group and between sex of the same group, with 95% confidence intervals. No significant difference detected.

**Table S1.**
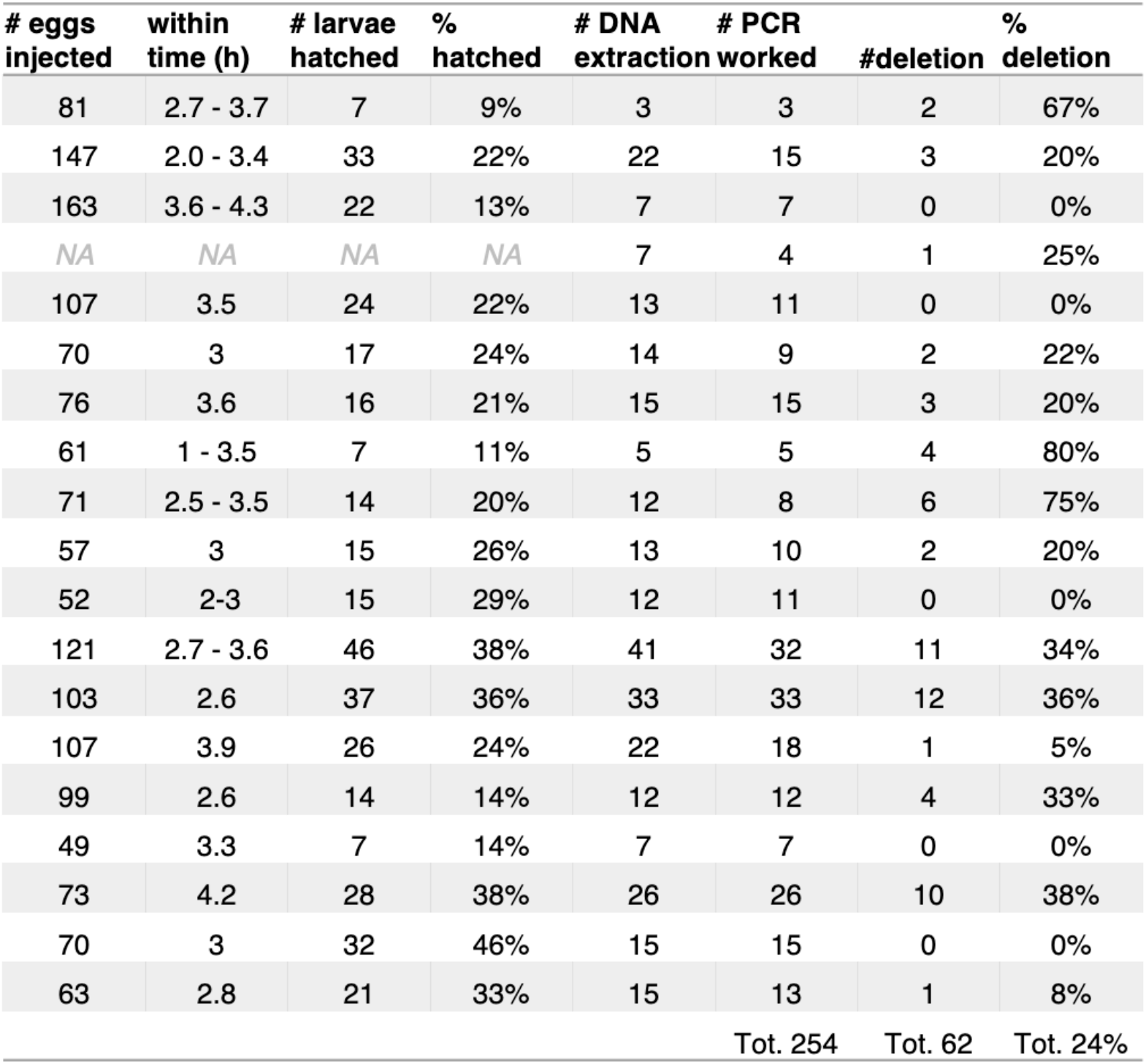
Survival and efficiency statistics in CRISPR/Cas9 experiments. In all the injections above the same concentration and mix of 2 sgRNAs targeting *regucalcin1* were used (see Table S2), with a sgRNA to Cas9 concentration of 250/500 ng/µl.

**Table S2.**
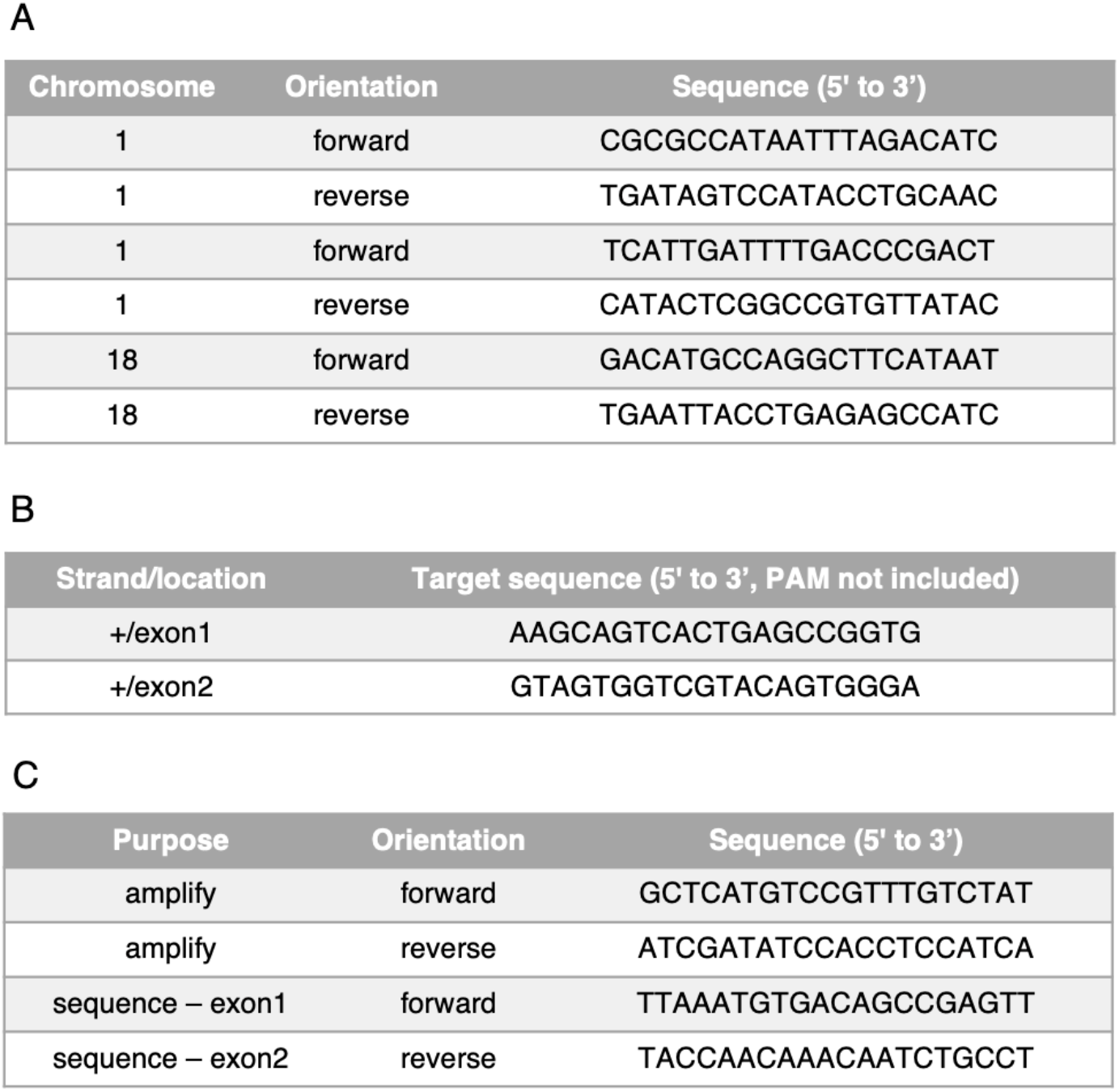
Primer and guide RNA sequences. (**A**) PCR primer sequences for obtaining genotype information at QTL locations (**B**) sgRNA sequences for CRISPR knock-outs of *regucalcin1* (**C**) PCR primer sequences for detecting *regucalcin1 KO* occurrence.

